# Heterologous expression of influenza hemagglutinin leads to early and transient activation of the unfolded protein response in *Nicotiana benthamiana*

**DOI:** 10.1101/2023.08.03.550084

**Authors:** Louis-Philippe Hamel, Marc-André Comeau, Rachel Tardif, Francis Poirier- Gravel, Marie-Ève Paré, Pierre-Olivier Lavoie, Marie-Claire Goulet, Dominique Michaud, Marc-André D’Aoust

## Abstract

The unfolded protein response (UPR) allows cells to cope with endoplasmic reticulum (ER) stress induced by the accumulation of misfolded proteins in the ER. Due to its sensitivity to *Agrobacterium tumefaciens*, model plant *Nicotiana benthamiana* is widely employed for the transient expression of recombinant proteins of biopharmaceutical interest, including therapeutic antibodies and virus surface proteins used for vaccine production. As such, study of the plant UPR is of practical significance, since enforced expression of complex secreted proteins often results in ER stress. After 6 days of expression, we recently reported that influenza hemagglutinin (HA) induces accumulation of UPR proteins. Since the upregulation of corresponding UPR genes was not detected at this time point, accumulation of UPR proteins was hypothesized to either be independent of transcriptional regulation, or associated with early but transient UPR gene upregulation. Using time course sampling, we here show that HA expression does result in early and transient activation of the UPR, as inferred from unconventional splicing of *NbbZIP60* transcripts and induction of UPR genes with varied functions. The transient nature of HA-induced UPR suggests that this response was sufficient to cope with ER stress provoked by expression of the secreted protein, as opposed to an antibody that triggered a stronger and more sustained UPR. As defense-related genes were induced after the peak of UPR activation and correlated with high increase in HA protein accumulation, we hypothesize that these immune responses, rather than the UPR, were responsible for the onset of necrotic symptoms on HA-expressing leaves.

**One-sentence summary:** *Agrobacterium*-mediated expression of influenza hemagglutinin results in early and transient activation of the unfolded protein response, preventing deleterious effects caused by unresolved endoplasmic reticulum stress.

## Introduction

In eukaryotes, many proteins rely on the cell secretory pathway for proper maturation, targeting and biological functions (Benham, 2012). These include secreted proteins, plasma membrane (PM) proteins and proteins targeted to lysosomes (and the vacuole in plant cells). To enter the secretory pathway, proteins are first translated on membrane-bound ribosomes located on the external face of the ER. Nascent polypeptides then reach luminal space of the ER via translocons, which act as protein channels for nascent protein translocation through the ER membrane. Inside the ER lumen, the so-called "client" proteins are rapidly subjected to chaperone-assisted folding and, when required, to post-translational modifications such as disulfide bridge formation, glycosylation, or oligomer assembly. To coordinate these activities, eukaryotic cells evolved a sophisticated protein machinery known as the ER quality control (ERQC) system (Araki and Nagata, 2011). While protein disulfide isomerases (PDIs) catalyse the formation of disulfide bonds, ER-resident chaperones of the Binding immunoglobulin Protein (BiP) family bind to exposed hydrophobic regions of unfolded proteins, helping them to adopt a proper tertiary structure. For glycoproteins, folding assistance is further provided by ER-resident lectins of the calreticulin (CRT) and calnexin (CNX) families.

ERQC allows for appropriate folding and modification of most client proteins, which in turn proceed towards downstream steps of the secretory pathway for further maturation, proper targeting, or secretion. Despite ERQC efficacy, some client proteins still fail to fold or assemble properly. When accumulation of misfolded proteins reaches a certain threshold, a second mechanism known as the ER-associated degradation (ERAD) system is activated (Araki and Nagata, 2011). ERAD allows for the retro-transport of misfolded proteins from the ER lumen to the cytosol, where degradation through the 26S proteasome will intervene. Distinct ERAD pathways exist (Brodsky and Wojcikiewicz, 2009; Wu and Rapoport, 2018), however the removal of soluble and ER membrane glycoproteins is mediated by luminal lectin osteosarcoma 9 (OS9; Hüttner *et al.,* 2012), which recognizes trimmed N-glycan chains of glycoproteins that have failed to fold properly (Liu and Howell, 2010). Together with the ER chaperone glucose-regulated protein 94 (GRP94), OS9 brings misfolded proteins to the retrotranslocation complex, which comprises ER membrane proteins such as Sel1L, HRD1, and Derlins (DERs). The retrotranslocation complex also comprises ATPase motor protein CDC48, which extracts misfolded proteins from the ER lumen and releases them in the cytosol. The E3 ubiquitin ligase activity of HRD1 then conjugates ubiquitin moieties to the target protein, marking it for degradation.

While ERQC and ERAD constantly maintain cell homeostasis, changes in environmental conditions or cell physiological status increase the needs for protein secretion. These changes can also induce conditions that are no longer favorable for ER protein folding and maturation. When accumulation of misfolded proteins overwhelms basal ERQC and ERAD functions, cells start to experience ER stress. To cope with ER stress, eukaryotes evolved refined signaling networks collectively known as the unfolded protein response (UPR; Read and Schröder, 2021). In plants, the UPR comprises two branches with conserved components and activation mechanisms (Liu and Howell, 2010; Iwata and Koizumi, 2012; Duwi Fanata *et al.,* 2013; Howell, 2013; 2021). In the first branch, membrane-tethered transcription factors (TFs) basic leucine zipper 17 (bZIP17) and bZIP28 are released from membrane anchoring via clipping of their transmembrane domain (TMD), an action mediated by proteases associated to the Golgi apparatus (Liu *et al.,* 2007). In the second branch, ER transmembrane sensor protein inositol requiring enzyme 1 (IRE1) uses its ribonuclease activity to unconventionally splice *bZIP60* transcripts, producing a shorter TF that lacks a TMD and therefore is no longer restrained by ER membrane anchoring (Nagashima *et al.,* 2011). Once activated, the UPR triggers a series of complementary mechanisms, including the shutdown of translational activities and the upregulation of UPR genes. These include ERQC components such *PDIs*, *BiPs*, *CNXs*, and *CRTs*, as well as genes encoding ERAD components (Kamauchi *et al.,* 2005; Iwata *et al.,* 2008; 2010). Purpose of the UPR is to restore cell homeostasis by reducing translation on one hand, and to increase ERQC and ERAD capabilities on the other hand. In case of severe ER stress, sustained activation of the UPR can also lead to the activation of programmed cell death, a mechanism that ultimately protects highly stressed tissues from cells that have become dysfunctional (Kørner *et al.,* 2015).

Plant molecular farming collectively refers to the approaches that make use of plant cells as biofactories to produce recombinant proteins or metabolites of pharmaceutical interest (Chung *et al.,* 2022). Recombinant proteins commonly produced *in planta* include therapeutic antibodies and surface proteins from mammalian viruses that are used for the production of vaccines. Using *Nicotiana benthamiana* leaf cells, the biopharmaceutical company Medicago for instance developed a molecular farming approach to produce influenza vaccine candidates at large-scale (D’Aoust *et al.,* 2008; Landry *et al.,* 2010). Based on bacterial vector *Agrobacterium tumefaciens*, this process relies on the transient expression of recombinant influenza hemagglutinins (HAs), along the viral suppressor of RNA silencing P19 (Silhavy *et al.,* 2002), which prevents silencing of recombinant *HA* genes delivered by the bacterium. Engineered to efficiently enter the ER, newly synthesized HA proteins travel through the plant cell secretory pathway, before being trafficked to the PM (D’Aoust *et al.,* 2008). When sufficient HA proteins have accumulated, bending of the PM allows for budding of the so-called virus-like particles (VLPs). These nanoscale assemblies comprise trimer clusters of the engineered HA protein embedded in a lipid envelope derived from the PM of plant cells. Structurally, VLPs and influenza viruses share similar size and shape, however the former lack genetic components required for replication. Once purified and formulated into vaccine candidates, VLPs induce an immune response that protects newly immunized hosts from subsequent infection by the influenza virus (Landry *et al.,* 2010).

We recently reported that HA protein accumulation results in a unique molecular signature that modifies metabolism and fitness of plant cells at 6 days post-infiltration (DPI), including the shutdown of chloroplast gene expression and the activation of plant immunity (Hamel *et al.,* 2023). A proteomics assessment at 6 DPI also revealed that HA protein expression results in the accumulation of UPR proteins, including PDIs, BiPs, and CRTs. Enforced expression of a complex secreted protein such as HA was expected to trigger the UPR so that plant cells can manage stress associated to the transient expression system, including increased needs for recombinant and endogenous defense protein secretion. Interestingly, thorough monitoring of the transcriptome at 6 DPI did not confirm induction of UPR genes, suggesting that increased accumulation of UPR proteins was either independent of transcriptional regulation, or that UPR gene upregulation occurred earlier before returning to levels preventing their detection at 6 DPI (Hamel *et al.,* 2023).

Using a time course sampling approach, we here show that HA protein expression does result in early and transient activation of the UPR, as inferred from the detection of unconventional splicing of *NbbZIP60* transcripts and the upregulation of UPR genes of various functions. The upregulation of UPR genes was detected at 3 DPI, but their expression had returned to basal level after 5 days. Our data also showed that activation of the UPR peaked prior to the induction of defense genes strongly responding to HA protein expression. Transient nature of the HA-induced UPR suggests that enhanced ERQC and ERAD functions were sufficient to cope with ER stress imposed by expression of the HA protein, as opposed to the expression of an antibody that induced a stronger and more sustained UPR. Overall, this work expands our understanding of host plant responses to foreign protein expression, in addition of providing the research community with a useful set of marker genes to study ER stress and associated UPR in *N. benthamiana*, a model plant and host of choice for molecular farming and study of plant immunity (Goodin *et al.,* 2008; Bally *et al.,* 2018; Chung *et al.,* 2022). Conserved molecular mechanisms linked to activation of the UPR in *N. benthamiana* are further discussed.

## Results

### Accumulation of UPR proteins in response to HA expression

To better define molecular responses in *N. benthamiana* leaves expressing influenza VLPs, we previously conducted a proteomics survey using isobaric tags for relative and absolute quantitation (iTRAQ) labelling (Hamel *et al.,* 2023). This revealed enhanced accumulation of UPR proteins at 6 DPI, including PDIs, BiPs, and CRTs (Table S1). Since the UPR plays a central role in protein secretion, we further investigated regulation of this pathway during foreign protein expression. Search of the *N. benthamiana* genome identified 21 PDIs (NbPDIs) that clustered similarly compared to PDIs from model plant *Arabidopsis thaliana* (AtPDIs; Figure S1A). Of the seven NbPDIs identified by proteomics, all but NbPDI16 and NbPDI17 displayed a C-terminal “KDEL” motif (Table S1), which works as an ER retention signal (Munro and Pelham 1987). Interestingly, identified NbPDIs also matched with a subset of AtPDIs previously involved in the UPR (Lu and Christopher, 2008), including NbPDI16 and NbPDI17 that lack an ER retention signal (Figure S1A). Search of the *N. benthamiana* genome also identified seven BiPs (NbBiPs; Figure S1B), and nine ER lectins that clustered in two sub-types corresponding to CRTs (NbCRTs) and CNXs (NbCNXs; Figure S1C). For these ER-resident chaperones, clustering was again similar to corresponding homologs in Arabidopsis, emphasizing conservation of ERQC components between the two plant species. For all NbBiPs and NbCRTs identified by proteomics, a C-terminal “HDEL” motif was identified, again suggesting retention in the ER (Table S1).

Our investigation of the changes in the transcriptome and the proteome at 6 DPI had also revealed upregulation of cytosolic heat shock proteins (HSPs), a response seen while expressing P19 alone, but not after co-expression of P19 and HA proteins (Hamel *et al.,* 2023). One exception to this was the gene model Niben101Scf04331g09018, which encodes a molecular chaperone of the HSP90 family. Unlike cytosolic HSPs described above, protein product from this gene harbors a C-terminal “KDEL” motif that suggests ER localization. The protein also showed enhanced accumulation following both P19 expression, or co-expression of P19 and HA proteins (Table S1). To the best of our knowledge, this HSP90 has never been formally characterized in *N. benthamiana*. Protein sequence alignments however showed that it shares homology to Arabidopsis SHEPHERD (AtSHD; AT4G24190), an ER-resident chaperone involved in the folding of CLAVATA proteins to regulate meristem growth (Ishiguro *et al.,* 2002). Product of the Niben101Scf04331g09018 gene is also closely related to human GRP94, a key ERAD component that interacts with OS9 to deliver misfolded proteins to the retrotranslocation complex (Christianson *et al.,* 2008; Marzec *et al.,* 2012; Seidler *et al.,* 2014). Herein termed NbGRP94, over-accumulation of that ER chaperone further suggests that the UPR was activated during HA protein secretion. We hypothesized that this response contributes to efficient expression of the HA protein in addition of preventing negative effects of the ER stress perhaps induced during enforced expression of the secreted protein.

### Time course sampling, stress symptoms, and recombinant protein accumulation

To assess whether the accumulation of UPR proteins was dependant on early upregulation of the corresponding UPR genes, time course sampling was performed at 1, 3, 5, 7, and 9 DPI. For each time point, leaves from non-infiltrated (NI) plants, or plants only infiltrated with resuspension buffer (Mock) were used as controls. Sampling was also performed for agroinfiltrated leaves expressing P19 only, or co-expressing P19 along with the HA protein from pandemic influenza virus strain H5 Indonesia (H5/A/Indonesia/05/2005; H5^Indo^; Hamel *et al.,* 2023). Hereafter referred to as H5, this HA protein results in formation of VLPs *in planta*. Regulation of the UPR was also assessed during co-expression of P19 along with two monoclonal antibodies (mAbs), mAb1 and mAb2. Within plant cells, mAbs are produced by the secretion and assembly of antibody chain proteins, but these do not form VLPs. The two mAbs expressed were selected based on (1) the contrasted stress symptoms they induced on the host plants, and (2) their contrasting accumulation levels, either high or very low, in leaf tissue following heterologous expression (see below).

The effects of foreign protein expression were first characterized by macroscopic evaluation of the stress symptoms induced on representative leaves from each condition harvested at 9 DPI (Figure 1A). Using NI leaves as a baseline, no obvious effect was visible on Mock-treated leaves. For agroinfiltrated leaves expressing P19 only, yellowish discoloration typical of chlorosis was visible, but no sign of plant cell death was detected. For leaves co-expressing P19 and H5 proteins, chlorosis was more advanced and accompanied by diffused greyish necrotic flecking that spread uniformly throughout the leaf blade. For leaves co-expressing P19 and mAb1, chlorosis was observed but again no sign of plant cell death was denoted. On the opposite, advanced plant cell death was detected on leaves co-expressing P19 and mAb2. Notably, necrotic lesions found on these leaves were somewhat different from those seen on H5-expressing leaves, with well-delimited cell death lesions spanning the entire leaf width.

**Figure 1.**
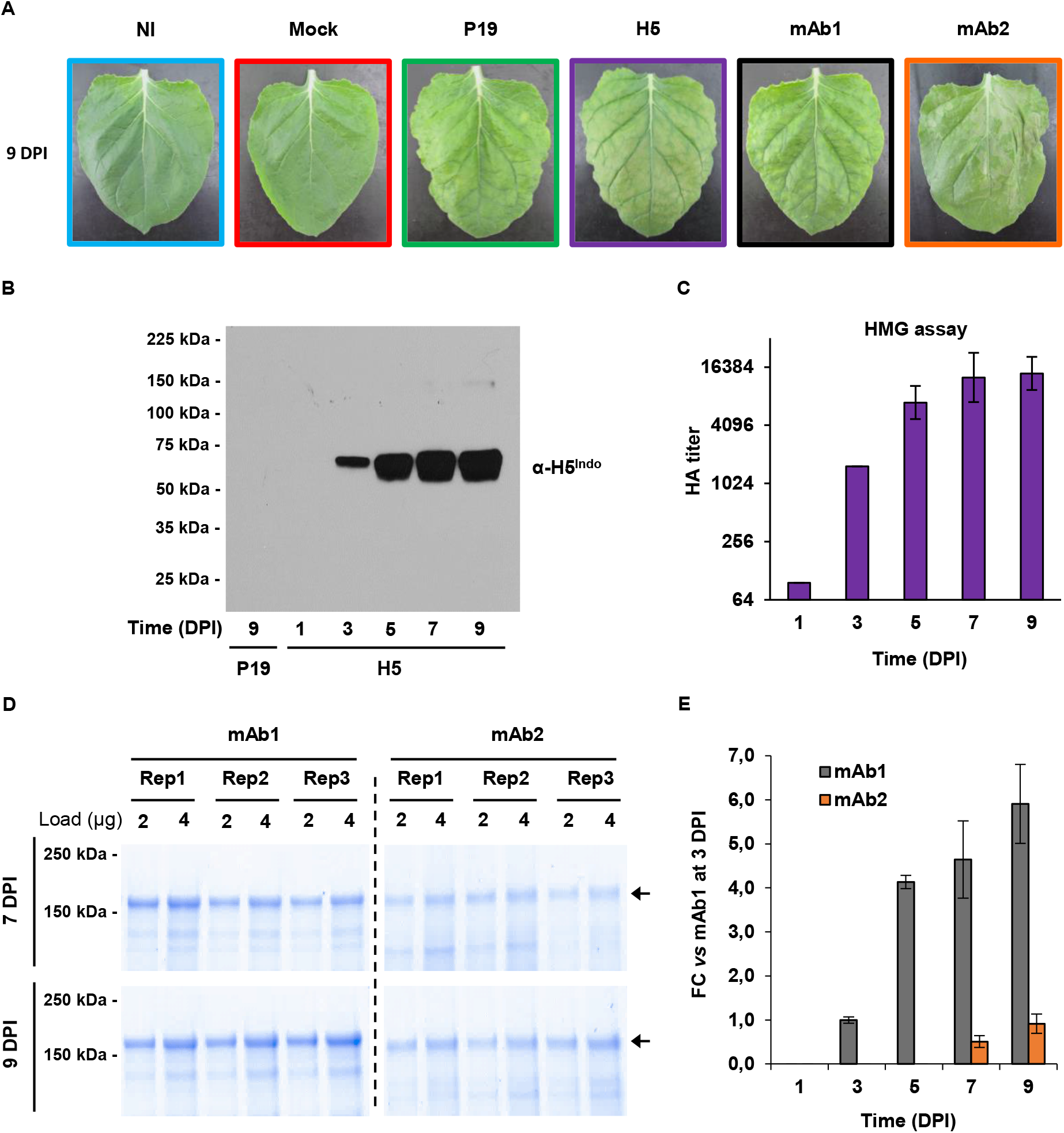
Stress symptoms and recombinant protein accumulation. (A) Stress symptoms observed on representative leaves from each condition harvested at 9 days post-infiltration (DPI). (B) Western blot depicting HA protein accumulation at each time point. A P19 sample harvested at 9 DPI was used as a control lacking HA protein expression. (C) Hemagglutination (HMG) assay depicting HA protein activity at each time point. (D) Quantification of complete monoclonal antibodies (mAb), as measured by densitometry (mAb1 on left panels and mAb2 on right panels). Accumulation measured at 7 and 9 DPI are shown (top and bottom panels, respectively). For each of three repetitions (Rep1 to Rep3), 2 and 4 μg of total soluble proteins were loaded. Arrows indicate complete IgGs. (E) Fold-change (FC) accumulation of mAbs, as measured by densitometry. Accumulation level of mAb1 at 3 DPI was arbitrarily set at 1-fold. Condition names are as follows: NI, non-infiltrated leaves; Mock, leaves infiltrated with buffer only; P19, leaves infiltrated with AGL1 and expressing P19 only; H5, leaves infiltrated with AGL1 and co-expressing P19 and H5^Indo^; mAb1 (and mAb2), leaves infiltrated with AGL1 and co-expressing P19, monoclonal antibody mAb1 (or mAb2) light (LC) and heavy (HC) chains.

To confirm the accumulation of H5 protein in leaves, a western blot analysis was performed using an antibody specific to the HA protein of influenza virus strain H5^Indo^ (Figure 1B). In H5 samples, no accumulation, and low accumulation of the HA protein were detected at 1 and 3 DPI, respectively. HA protein accumulation then increased notably between 3 and 5 DPI, and again between 5 and 7 DPI. At that point, H5 accumulation had pretty much reached a plateau, as the level observed at 9 DPI was roughly the same as the one seen at 7 DPI (Figure 1B). To assess HA activity, hemagglutination (HMG) assays were conducted (Figure 1C). At 1 DPI, HA activity was barely detectable, consistent with the absence of measurable H5 product (Figure 1B). HA activity then increased between 1 and 7 DPI to reach a plateau maintained up to 9 DPI (Figure 1C). Overall, H5 accumulation and HA activity correlated tightly, confirming that recombinant H5 accumulated *in planta* was still active against red blood cell receptors following extraction from leaf tissues.

To monitor the accumulation of recombinant mAb1 and mAb2, gel densitometry measurements were performed on assembled antibodies produced *in planta*. At 7 and 9 DPI, Coomassie blue-stained gels revealed higher accumulation of mAb1 compared to mAb2 (Figure 1D). By arbitrarily setting the accumulation of mAb1 at 3 DPI to one-fold, relative fold-change (FC) accumulation values were determined for each time point (Figure 1E). No accumulation of mAb1 was detected at 1 DPI, but levels significantly increased between 3 and 5 DPI, reaching more than four-fold accumulation level compared to the baseline. The mAb1 accumulation rate then decreased between 5 and 9 DPI, but the accumulation peak was not reached until the last leaf sampling point at 9 DPI. For mAb2, product accumulation could not be detected until 7 DPI (Figure 1E). mAb2 level then slightly increased between 7 and 9 DPI, but overall accumulation remained much lower compared to mAb1 (Figure 1E). Considering these poor accumulation levels and the strong stress symptoms induced by *in planta* expression of mAb2 (Figure 1A), we hypothesized that this product leads to an unresolved ER stress and therefore that regulation of UPR gene expression within these samples would be informative for comparison with P19, mAb1, and H5 samples.

### Activation of the IRE1-bZIP60 pathway

In the absence of ER stress, IRE1 is held in a monomeric, non-signaling state through interaction with luminal BiPs. When misfolded proteins start to accumulate, BiPs get recruited for protein folding, freeing the luminal domain of IRE1 that also interacts with misfolded proteins (Figure 2A). Following protein oligomerization and phosphorylation, IRE1 gets activated and using its cytosolic RNase domain, it performs unconventional splicing of *bZIP60* transcripts. The shorter TF produced lacks a TMD and as a result translocates to the nucleus where it upregulates expression of UPR genes that carry specific *cis* regulatory elements in their promoter (Figure 2A; Liu and Howell, 2016; Li and Howell, 2021).

**Figure 2.**
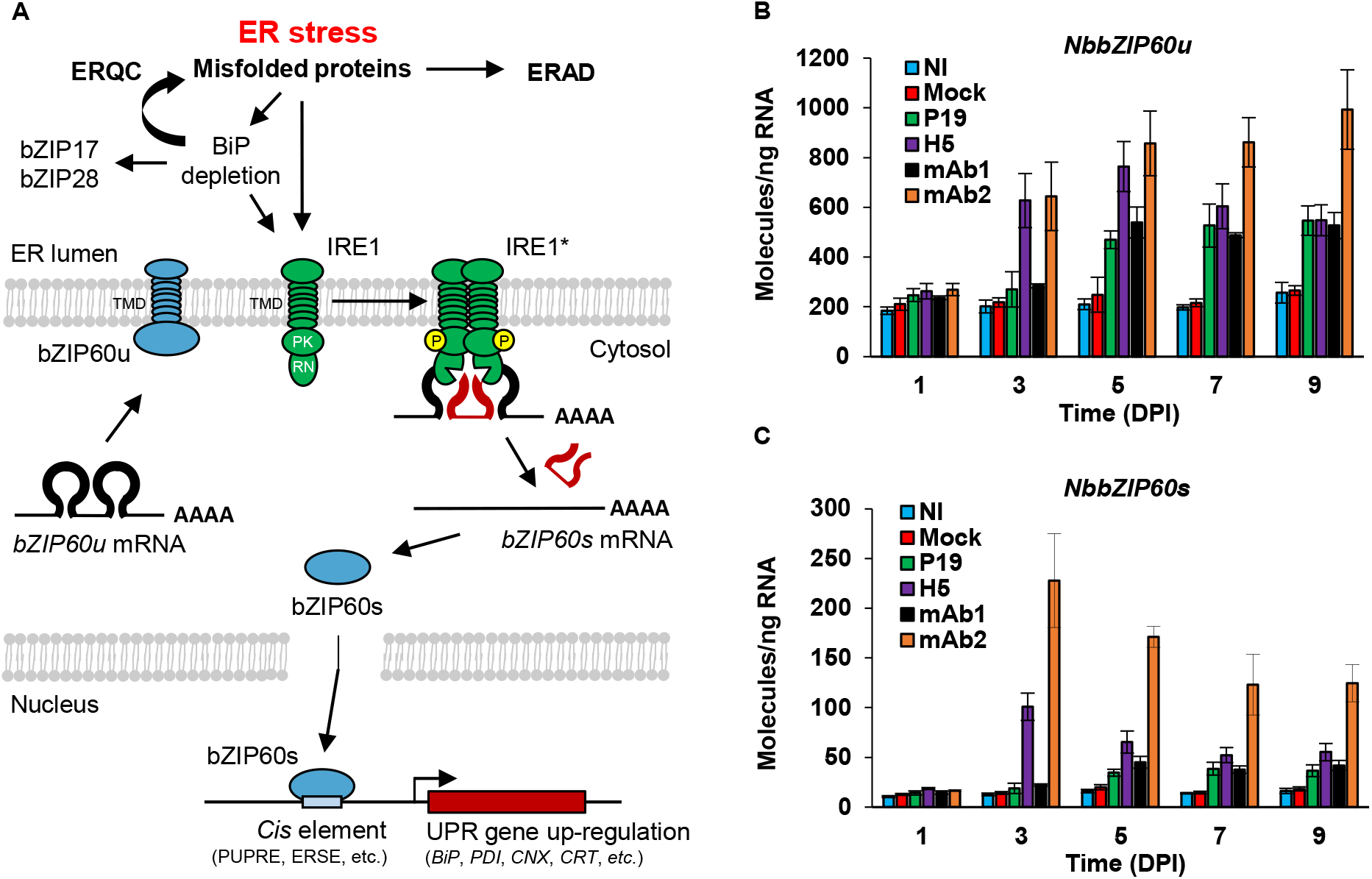
The IRE1-bZIP60 pathway and upregulation of *NbbZIP60*. (A) Model depicting unconventional splicing of *bZIP60* transcripts by IRE1, an ER transmembrane sensor that owns protein kinase (PK) and ribonuclease (RN) activity. In the absence of stress, unspliced *bZIP60* transcripts (*bZIP60u*) produce a transcription factor (TF) that harbors a transmembrane domain (TMD). Membrane tethering of bZIP60 prevents its translocation to the nucleus. ER stress results in misfolded protein accumulation and consequent ERQC and ERAD activation. This eventually triggers IRE1 activation (IRE1*) through oligomerization and phosphorylation (P). Unconventional splicing of *bZIP60* transcripts (*bZIP60s*) results in translation of a shorter TF that lacks a TMD and that is now free to induce UPR gene expression in the nucleus. Within the promoter of UPR genes, bZIP60s recognizes *cis* regulatory elements such as the plant unfolded protein response element (UPRE) or the ER stress-response element (ERSE). Adapted from Duwi Fanata *et al.,* 2013. To assess activation of the UPR, expression of *NbbZIP60u* (B) and of *NbbZIP60s* (C) was measured by RTqPCR. For each time point in days post-infiltration (DPI), results are expressed in numbers of molecules per ng of RNA. Condition names are as follows: NI, non-infiltrated leaves; Mock, leaves infiltrated with buffer only; P19, leaves infiltrated with AGL1 and expressing P19 only; H5, leaves infiltrated with AGL1 and co-expressing P19 and H5^Indo^; mAb1 (and mAb2), leaves infiltrated with AGL1 and co-expressing P19, monoclonal antibody mAb1 (or mAb2) light (LC) and heavy (HC) chains.

To investigate IRE1 activation and unconventional splicing of *bZIP60* transcripts, we retrieved nucleotide sequences of Niben101Scf24096g00018, the gene model encoding *N. benthamiana*’s version of the UPR-related TF bZIP60. Herein termed *NbbZIP60*, this gene displayed strict sequence conservation around predicted splicing sites recognized by IRE1 (Figure S2). Based on this sequence conservation, forward primers specific to the spliced (s) or unspliced (u) versions of *NbbZIP60* transcripts were designed (Figure S2). Paired to a reverse primer that recognized both transcripts, forward primers were used to selectively profile expression of *NbbZIP60u* and *NbbZIP60s* via real time quantitative polymerase chain reaction (RTqPCR). For *NbbZIP60u*, results showed similar expression levels for all conditions at 1 DPI (Figure 2B). For NI and Mock samples, expression levels of *NbbZIP60u* remained low and unchanged up to 9 DPI. For H5 and mAb2 samples, upregulation of *NbbZIP60u* was seen at 3 DPI and extent of the response was similar for both conditions. In H5 samples, expression of *NbbZIP60u* slightly increased between 3 and 5 DPI, before going down slightly at 7 and 9 DPI. For mAb2 samples, expression level of *NbbZIP60u* also increased between 3 and 5 DPI, but the expression levels then remained high up to the end of leaf sampling at 9 DPI. Compared to NI and Mock controls, upregulation of *NbbZIP60u* was also observed in P19 and mAb1 samples, but this response only arose later (5 DPI) and remained weaker compared to H5 and mAb2 samples (Figure 2B).

For *NbbZIP60s*, overall expression levels were lower than those observed for *NbbZIP60u* (see Y axis scales on Figures 2B and 2C). This suggests that only a fraction of *NbbZIP60* transcripts were unconventionally spliced by IRE1. At 1 DPI, very few spliced transcripts were detected for any of the conditions (Figure 2C), while significantly higher levels of *NbbZIP60s* transcripts were detected for H5 and mAb2 samples at 3 DPI (with significantly higher levels for the latter). For both conditions, levels of spliced transcripts then slowly decreased between 3 and 9 DPI, with levels from mAb2 samples again remaining significantly higher compared to other conditions, including H5 samples. At 5 DPI, H5 samples still displayed significantly higher levels of spliced transcripts compared to P19 samples, but this was no longer the case at 7 and 9 DPI (Figure 2C). For NI and Mock samples, levels of spliced transcripts remained low and unchanged throughout the time course. Overall, our data suggests early and transient activation of IRE1 in H5 samples, while early and more sustained activation of IRE1 was seen in mAb2 samples. Compared to NI and Mock controls, IRE1 splicing activity was also detected in P19 and mAb1 samples, although arising later and remaining weaker compared to H5 or mAb2 samples.

### Upregulation of ERQC genes

Accumulation of *NbbZIP60s* transcripts (Figure 2C) suggests the UPR to be activated with different strength and kinetics, depending on the foreign protein combinations expressed. Activation of the UPR generally leads to the upregulation of genes encoding ERQC components, including PDIs and ER-resident chaperones of the BiP, CRT, and CNX families (Kamauchi *et al.,* 2005; Iwata *et al.,* 2008; 2010). Based on proteomics results (Table S1) and homology with homologs from Arabidopsis (Figure S1), primers specific to selected ERQC genes were designed to perform RTqPCR. For NI and Mock samples, expression of ERQC genes remained low and essentially unchanged throughout expression (Figure 3A for *NbPDIs*, Figure 3B for *NbBiPs*, Figure S3A for *NbCRTs*, and Figure S3B for *NbCNXs*). For the other conditions, slight differences between respective gene expression profiles were identified, however three major trends emerged. An early and transient expression pattern was observed for H5 samples, characterized by an upregulation of ERQC genes at 3 DPI followed by expression levels rapidly decreasing to reach levels similar to those observed in P19, mAb1, or even NI and Mock control samples at 5 DPI. A second expression pattern was observed in mAb2 samples, also characterized by early an upregulation of ERQC genes at 3 DPI. Unlike the first pattern, ERQC gene expression was however sustained at later time points, with levels that never dropped back to basal level and remained significantly higher compared to the other conditions. The third expression pattern was observed in P19 and mAb1 samples, characterized by a weaker upregulation of ERQC genes at 3 DPI. At later time points, the expression of ERQC genes was above the expression levels observed for NI and Mock controls, but always remained significantly lower compared to H5 and mAb2 samples. Overall, these observations at 3 DPI suggest that the UPR was more strongly activated in H5 and mAb2 samples than in other conditions. For some ERQC genes, they also suggest a higher response level in mAb2 samples than in H5 samples (e.g. *NbBiP2a/b* and *NbBiP3a/b* on Figure 3B). Expression patterns from most ERQC genes examined also tightly correlated with accumulation patterns of *NbbZIP60s* transcripts (Figure 2C), suggesting that these UPR genes are direct genetic targets of the TF.

**Figure 3.**
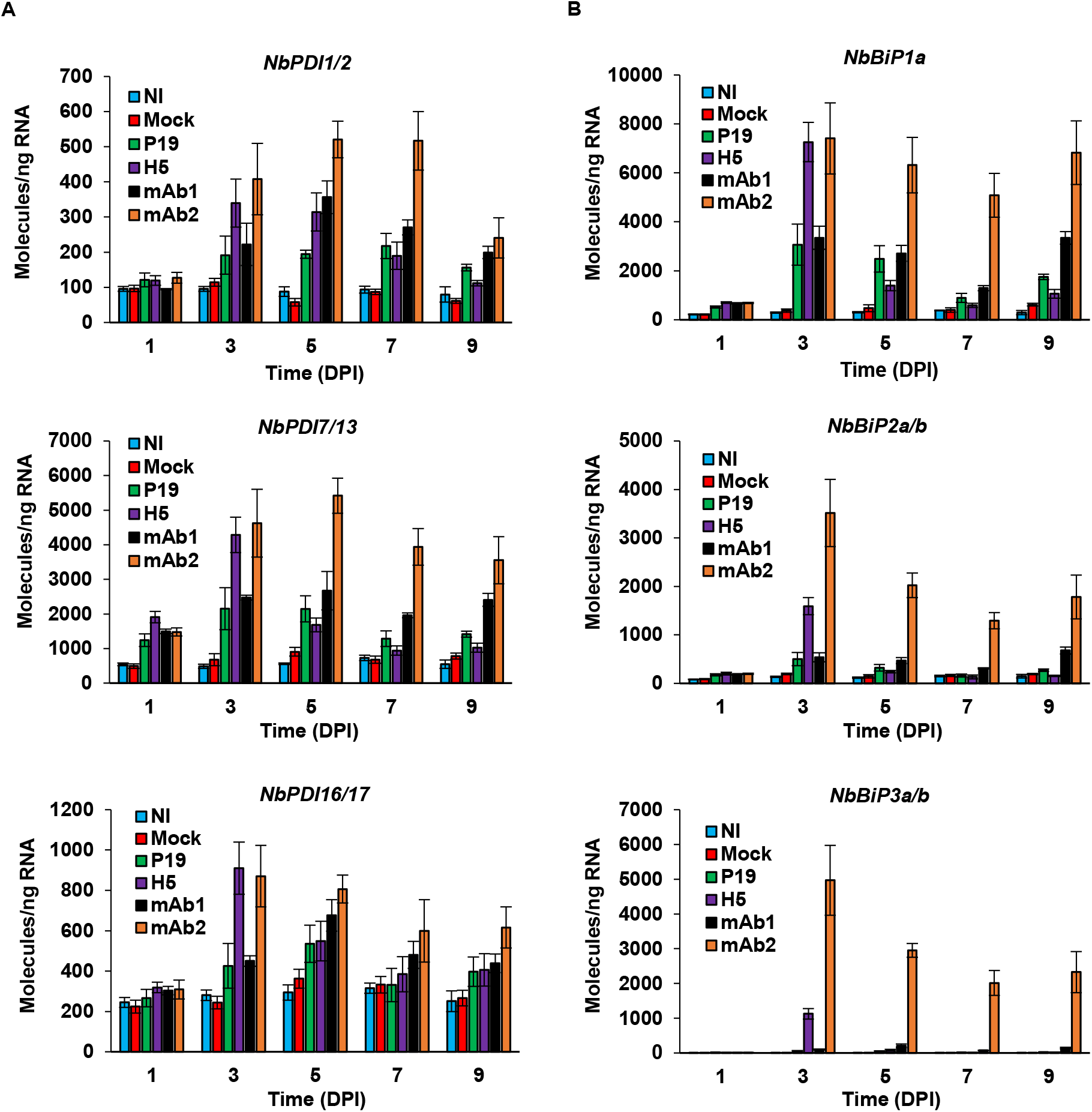
Expression of ERQC genes: *PDIs* and *BiPs*. Expression of genes encoding for PDIs (A) or ER resident chaperones of the BiP family (B), as measured by RTqPCR. For each time point in days post-infiltration (DPI), results are expressed in numbers of molecules per ng of RNA. Condition names are as follows: NI, non-infiltrated leaves; Mock, leaves infiltrated with buffer only; P19, leaves infiltrated with AGL1 and expressing P19 only; H5, leaves infiltrated with AGL1 and co-expressing P19 and H5^Indo^; mAb1 (and mAb2), leaves infiltrated with AGL1 and co-expressing P19, monoclonal antibody mAb1 (or mAb2) light (LC) and heavy (HC) chains.

### Upregulation of ERAD genes

In Arabidopsis, activation of the UPR also promotes the expression of ERAD genes (Kamauchi *et al.,* 2005; Iwata *et al.,* 2008; 2010). These include *Sel1L*, *HRD1*, *DER2*, and *DER3*, which all encode components of the retrotranslocation complex (Figure 4A). To study ERAD gene expression during foreign protein accumulation, we searched the genome of *N. benthamiana* to identify ERAD gene homologs and profiled expression for some of the retrieved candidates using RTqPCR. Selected genes were *NbSel1La* and *NbSel1Lb* (Figure 4B), *NbHRD1a* (Figure 4C), as well as closely related *NbDER2* and *NbDER3* (Figure 4D). At 1 DPI, the respective expression levels of all these genes were similar for all conditions. At 3 DPI, ERAD gene expression significantly increased to similar levels in H5 and mAb2 samples, while P19 and mAb1 samples had similar expression levels compared to NI and Mock controls. At 5 DPI, ERAD gene expression from H5 samples had decreased importantly to reach levels that were similar to P19 and mAb1 samples. After 3 DPI, ERAD gene expression from mAb2 samples also tended to decrease, but always remained significantly higher compared to other conditions. Overall expression patterns from ERAD genes examined were somewhat similar to those seen above for *NbbZIP60s* (Figure 2C) and ERQC genes (Figures 3 and S3).

**Figure 4.**
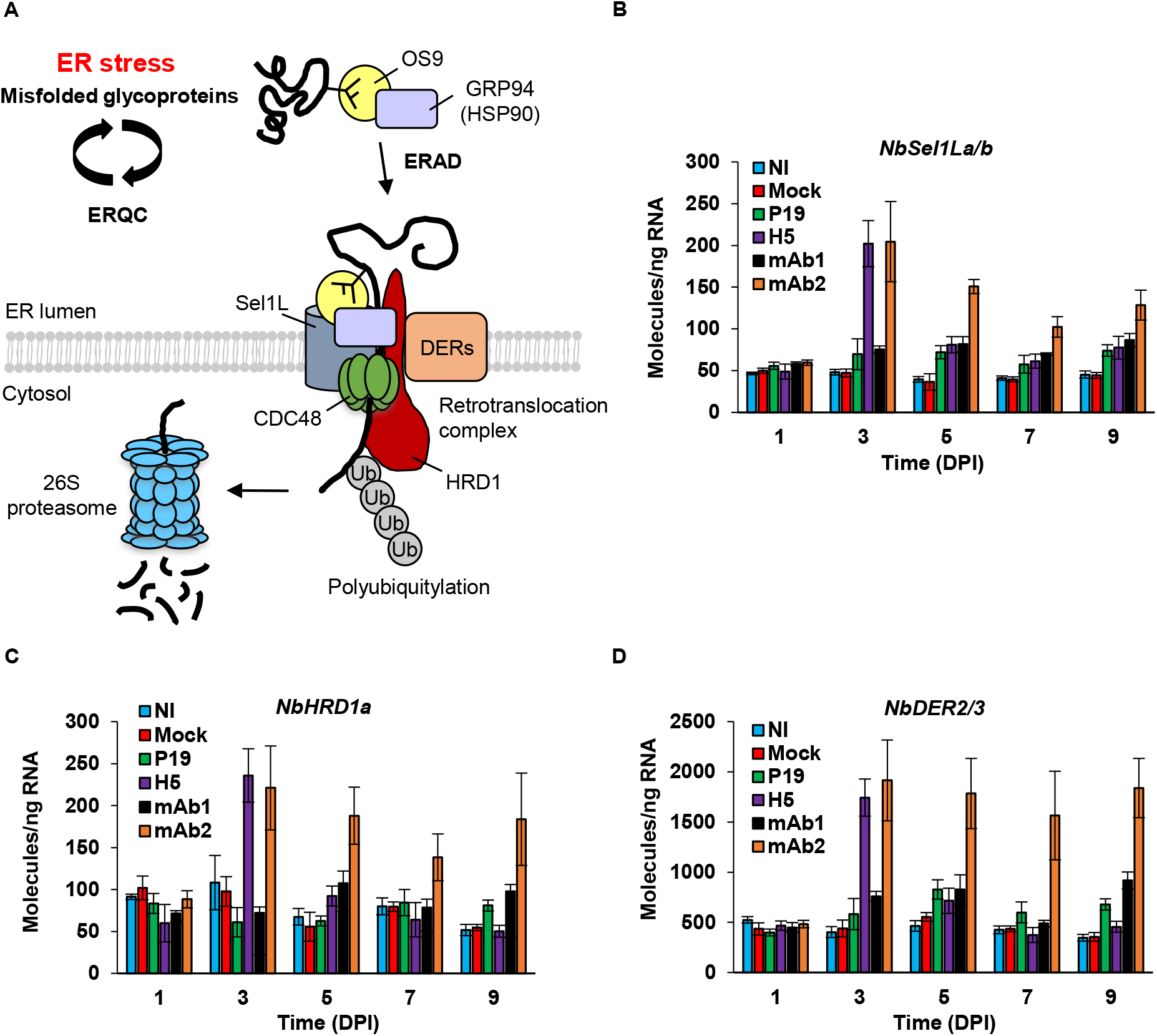
The ERAD system and expression of ERAD genes. (A) Model depicting ER retrotranslocation and cytosolic degradation of a soluble glycoprotein that was irrevocably misfolded. After trimming of its N-glycan chains, the glycoprotein is recognized by luminal lectin OS9. Together with GRP94, a molecular chaperone of the HSP90 family, OS9 brings its target to the retrotranslocation complex. The latter comprises ER membrane proteins Sel1L, HRD1, and Derlins (DERs). Through ATPase motor CDC48, the misfolded glycoprotein is extracted from the ER lumen and released in the cytosol. E3 ubiquitin ligase activity of HRD1 then catalyses conjugation of ubiquitin (Ub) moieties to the misfolded glycoprotein, targeting it for degradation via the 26S proteasome. Adapted from Howell, 2021. Expression of genes encoding for Sel1L (B), HRD1 (C), or DER (D) proteins, as measured by RTqPCR. For each time point in days post-infiltration (DPI), results are expressed in numbers of molecules per ng of RNA. Condition names are as follows: NI, non-infiltrated leaves; Mock, leaves infiltrated with buffer only; P19, leaves infiltrated with AGL1 and expressing P19 only; H5, leaves infiltrated with AGL1 and co-expressing P19 and H5^Indo^; mAb1 (and mAb2), leaves infiltrated with AGL1 and co-expressing P19, monoclonal antibody mAb1 (or mAb2) light (LC) and heavy (HC) chains.

### Other UPR genes: subunits of the Sec61 translocon and Bax inhibitor-1

Translocons are protein complexes that transport other proteins from their translation site in the cytosol to the ER lumen. These structures comprise several subunits, including the heterotrimeric complex Sec61 that forms the translocon pore and is made from assembly of secretory (Sec) proteins Sec61α, Sec61β, and Sec61γ (Itskanov and Park, 2023). As a way to increase protein secretion under stress conditions, genes encoding Sec proteins are induced during the UPR (Kamauchi *et al.,* 2005; Iwata *et al.,* 2008; 2010). Search of the *N. benthamiana* genome identified three *Sec61*α genes, six *Sec61*β genes, and five almost identical *Sec61*γ genes (Figure S4A). To examine expression of some of these candidates using RTqPCR, primers were designed to target *NbSec61*α*-1* (Figure S4B) or closely related *NbSec61*β*-1* and *NbSec61*β*-2* (Figure S4C). Similar to ERQC and ERAD genes, results showed an enhanced transcript accumulation that started at 3 DPI, a response more importantly induced in H5 and mAb2 samples compared to P19 and mAb1 samples. For mAb2 samples, *NbSec61* gene expression remained high until the end of sampling at 9 DPI, while it rapidly decreased in H5 samples at 5 DPI to reach levels similar to those observed in P19 and mAb1 samples. No gene induction was seen in NI and Mock controls.

*Bax inhibitor-1* (*BI-1*) genes encode ER transmembrane proteins first identified as suppressors of the cell death induced by pro-apoptotic protein Bax from yeast and mammals (Hückelhoven, 2004). In the case of severe ER stress, BI-1 protein levels act as a rheostat controlling activation of programmed cell death, including in plants (Ishikawa *et al.,* 2011). This function perhaps explains why *BI-1* genes are induced during the plant UPR (Kamauchi *et al.,* 2005; Iwata *et al.,* 2008; 2010). Search of the *N. benthamiana* genome identified at least four BI-1 homologs (NbBI-1s), which were most closely related to AtBI-1a of Arabidopsis (AT5G47120; Figure S4D). Primers specific to *NbBI-1a* and *NbBI-1b* showed these genes to be induced at 3 DPI in all agroinfiltrated conditions compared to NI and Mock controls. However, upregulation levels were significantly higher in H5 and mAb2 samples compared to P19 and mAb1 samples (Figure S4E). At later time points, higher expression seen in H5 samples reverted to levels similar to those seen in P19 and mAb1 samples, while expression remained significantly higher in mAb2 samples. Overall, these results confirmed that the expression of UPR genes of varied functions was induced in H5 and mAb2 samples, and again that these expression profiles mirrored the expression profile of *NbbZIP60s* (Figure 2C).

### Kinetics of defense gene expression vs activation of the UPR

Expression of the H5 protein was previously shown to induce a number of plant immune responses at 6 DPI, including the upregulation of oxylipin regulatory and response genes (Hamel *et al.,* 2023). As these observations were made at a single time point, we used RTqPCR to profile expression of some of these defense genes throughout the expression phase and compared regulation of immune responses with activation timeline of the UPR. For oxylipin regulatory genes, we monitored expression of patatin gene *NbPAT1*, of 9-lipoxygenase gene *NbLOX1*, of divinyl ether synthase (DVE) or epoxy alcohol synthase (EAS) genes *NbCYP74a* and *NbCYP74b*, and of allene oxide synthase gene *NbAOS1* (Hamel *et al.,* 2023). For *NbPAT1* (Figure 5A), *NbLOX1* (Figure 5B), and *NbCYP74* genes (Figure 5C), enhanced gene expression was mostly seen in H5 and mAb2 samples, with higher expression levels detected in the former case. Upregulation of these genes also arose faster in H5 samples, with significantly enhanced expression detected at 3 DPI. To a much lesser extent, upregulation of these genes was also detected in P19 and mAb1 samples compared to NI and Mock controls. This was consistent with the reported low induction of these genes by *Agrobacterium* (Hamel *et al.,* 2023). For *NbAOS1*, a weak upregulation was detected in mAb2 samples at 7 and 9 DPI, but gene induction again started earlier (5 DPI) and reached much higher levels in H5 samples (Figure 5D).

**Figure 5.**
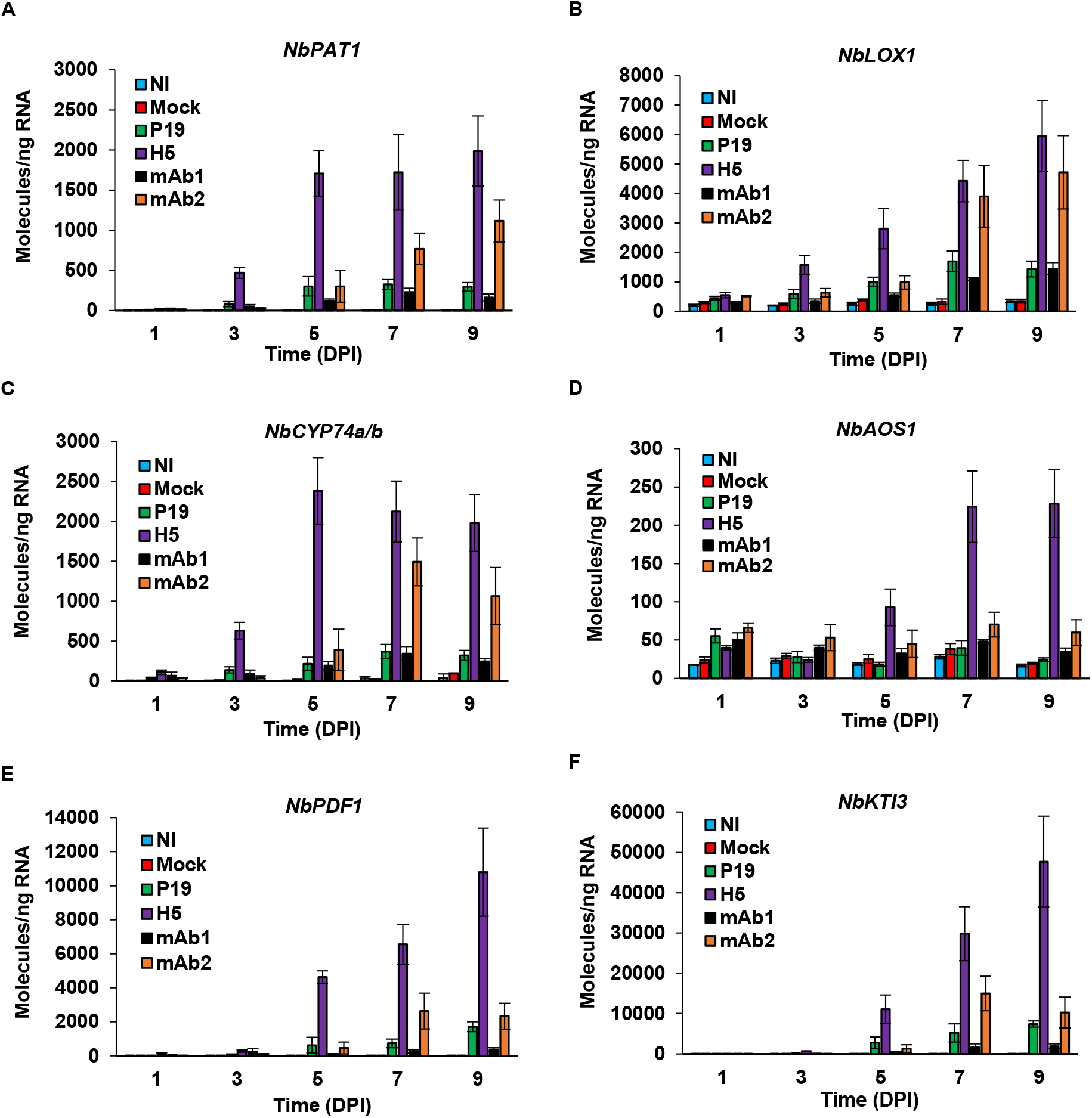
Expression of oxylipin-related genes. Expression of oxylipin regulatory genes *NbPAT1* (A), *NbLOX1* (B), *NbCYP74a* and *NbCYP74b* (C), as well as *NbAOS1* (D), as measured by RTqPCR. The expression of oxylipin response genes *NbPDF1* (E) and *NbKTI3* (F) was also assessed. These genes were selected because of their responsiveness to HA protein expression (Hamel *et al.,* 2023). For each time point in days post-infiltration (DPI), results are expressed in numbers of molecules per ng of RNA. Condition names are as follows: NI, non-infiltrated leaves; Mock, leaves infiltrated with buffer only; P19, leaves infiltrated with AGL1 and expressing P19 only; H5, leaves infiltrated with AGL1 and co-expressing P19 and H5^Indo^; mAb1 (and mAb2), leaves infiltrated with AGL1 and co-expressing P19, monoclonal antibody mAb1 (or mAb2) light (LC) and heavy (HC) chains.

For oxylipin response genes, expression patterns of plant defensin gene *NbPDF1* (Figure 5E) and of Kunitz trypsin inhibitor gene *NbKTI3* (Figure 5F) were examined. These genes were previously shown to be highly responsive to HA protein expression (Hamel *et al.,* 2023). Consistent with the induction of oxylipin regulatory genes, RTqPCR revealed strongest upregulation in H5 samples. In both cases, enhanced expression was not detected at 3 DPI, but was obvious at 5 DPI (Figure 5E and Figure 5F). Expression levels from both genes then continued to increase until the end of sampling at 9 DPI. At 7 and 9 DPI, the two genes were also upregulated in P19 and mAb2 samples, but their expression levels were significantly lower than in H5 samples. Together, these results indicate that upregulation of oxylipin regulatory genes preceded upregulation of oxylipin response genes. For the H5 samples, results also indicate that activation of the oxylipin pathway occurred after activation of the UPR, which peaked at 3 DPI in this condition.

At 6 DPI, transcriptomics analyses had also shown strong, and in many cases, H5-specific induction of genes promoting the activation of oxidative stress responses, including in the apoplast where VLP accumulation takes place (Hamel *et al.,* 2023). In view of this, we profiled the expression of some of these genes, namely NADPH oxidase gene *NbRBOHd*, closely related polyphenol oxidase genes *NbPPO1* and *NbPPO3*, secreted carbohydrate oxidase gene *NbBBE2*, as well as secreted ascorbate oxidase genes *NbAO1* and *NbAO2* (Hamel *et al.,* 2023). Results at 1 DPI indicated a similar upregulation of *NbRBOHd* in Mock and agroinfiltrated samples compared to NI samples (Figure 6A). A comparable expression pattern was observed at 3 DPI but not at 5 DPI, as *NbRBOHd* was expressed at significantly higher level in H5 samples compared to other agroinfiltrated conditions. Expression of the gene in H5 samples decreased at 7 and 9 DPI, but still remained significantly higher compared to P19, mAb1, or mAb2 samples. For *NbPPO* genes (Figure 6B) and *NbBBE2* (Figure 6C), RTqPCR revealed similar expression profiles, with stronger upregulation again seen in H5 samples. In this latter case, upregulation was first detected at 5 DPI and then steadily increased until the end of sampling at 9 DPI. For the last two time points, enhanced gene expression was also detected in mAb2 samples and, to a lower extent, in P19 samples. In both cases, defense gene expression however remained significantly lower compared to H5 samples. No or very limited expression of these genes was detected in NI, Mock, or mAb1 samples. For *NbAO* genes (Figure 6D), a significant induction was detected in all agroinfiltrated leaf samples compared to NI and Mock controls. In all cases, the highest upregulation level was reached at 5 DPI, even though gene induction started earlier and reached significantly higher levels in H5 samples. Taken together, these results suggest that oxidative stress signaling was stronger in H5 and mAb2 samples, correlating with cell death symptoms observed on leaves expressing these products (Figure 1A). For H5-expressing samples, upregulation of oxidative stress-related genes again came after the peak of UPR activation at 3 DPI.

**Figure 6.**
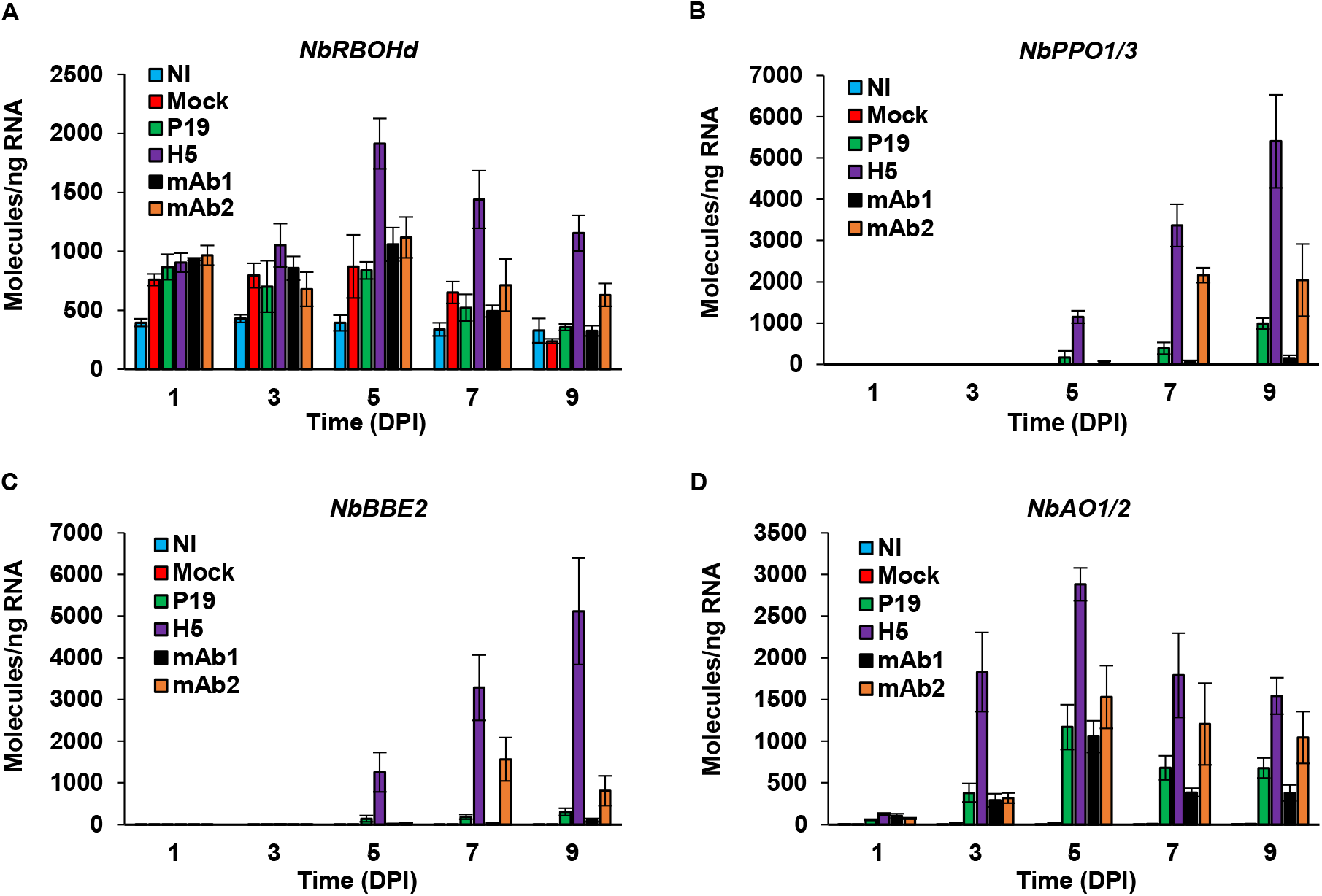
Expression of oxidative stress-related genes. Expression of *NbRBOHd* (A), *NbPPO1* and *NbPPO3* (B), *NbBBE2* (C), as well as *NbAO1* and *NbAO2* (D), as measured by RTqPCR. These genes were selected because of their responsiveness to HA protein expression (Hamel *et al.,* 2023). For each time point in days post-infiltration (DPI), results are expressed in numbers of molecules per ng of RNA. Condition names are as follows: NI, non-infiltrated leaves; Mock, leaves infiltrated with buffer only; P19, leaves infiltrated with AGL1 and expressing P19 only; H5, leaves infiltrated with AGL1 and co-expressing P19 and H5^Indo^; mAb1 (and mAb2), leaves infiltrated with AGL1 and co-expressing P19, monoclonal antibody mAb1 (or mAb2) light (LC) and heavy (HC) chains.

## Discussion

Plant molecular farming allows for the large-scale production of many clinically useful proteins, including therapeutic antibodies and virus surface proteins such as influenza HAs. As such, plant biotechnology offers alternative approaches to prevent the spreading of infectious diseases and to limit the societal and economic burdens associated with these diseases (Ortiz de Lejarazu-Leonardo *et al.,* 2021). Despite promising developments, mass infiltration of *Agrobacterium* and high yield production of foreign proteins unavoidably put pressure on host plants, including the cellular machinery mediating protein folding and maturation in the ER. Understanding the impact of those stresses is an important step to ensure sustainability of molecular farming approaches, especially since proteins with biopharmaceutical interest generally require refined protein domain folding and post-translational modifications such as sophisticated glycosylation patterns or assembly in quaternary conformations. Overloading the host cell ER with such complex proteins can lead to ER stress and, in turn, in activation of the UPR. When unresolved, ER stress can dramatically compromise plant cell fitness and viability, resulting in poor biomass quality and insufficient yields at harvest.

### Protein-specific activation of the UPR

At 6 DPI, iTRAQ proteomics revealed that *Agrobacterium*-mediated expression of influenza protein H5 induces accumulation of UPR proteins of various functions (Table S1; Hamel *et al.,* 2023). Interestingly, transcriptomics did not confirm an upregulation of the corresponding UPR genes at 6 DPI, suggesting this response to be independent of UPR gene induction, or to involve early and transient upregulation of these genes prior to leaf biomass harvesting. Here, time course sampling showed that HA protein expression does in fact result in the induction of UPR genes, a response initiated early following agroinfiltration between 1 and 3 DPI. UPR gene upregulation also correlated tightly with unconventional splicing of *NbbZIP60* transcripts, the closest homolog of *bZIP60* that controls activation of the UPR in Arabidopsis (Iwata and Koizumi, 2005; Iwata *et al.,* 2008). Since unconventional splicing of *bZIP60* transcripts reflects the activation status of IRE1, this process is a reliable marker of UPR activation. Consistent with the lack of UPR gene induction at 6 DPI (Hamel *et al.,* 2023), H5-mediated activation of the UPR was transient and could no longer be detected after 5 days of heterologous protein expression.

In our attempt to better define importance of the UPR during foreign protein expression in *N. benthamiana*, time course sampling of mAb1- and mAb2-expressing leaves was also informative as the expression profiles obtained with these foreign protein combinations revealed distinct UPR activation patterns. Whereas mAb2 expression induced early, strong, and sustained activation of the UPR, expression of mAb1 resulted in late and much weaker activation of this pathway, with measured levels comparable to those induced by the expression of P19 only. Consistent with this, mAb2 resulted in heavy necrotic symptoms and poor yields, while the mAb1 product had little impact on biomass quality despite high product accumulation. Together, these results suggest that despite structural similarity between the two antibody products, expression of mAb2 led to an unresolved ER stress perhaps induced by improper folding of the antibody variable regions, or by unsuccessful quaternary protein structure assembly of antibody chains. In any case, these contrasted outcomes highlight the importance of UPR molecular components to guarantee success and sustainability of plant molecular farming approaches.

Considering the contrasted UPR activation patterns in mAb1 and mAb2 samples, we conclude that HA protein expression may have resulted in moderate activation of the UPR, at least for the H5^Indo^ viral strain used in this study. While the induction of UPR genes was obviously higher in H5 samples than in P19 or mAb1 samples, it was similar or lower than in mAb2 samples, depending on which UPR gene is considered. As for timing, initiation of the UPR occurred within the same timeframe in both H5 and mAb2 samples. The major difference between these two conditions therefore lied in duration of the response, which was transient in H5 samples and more sustained in mAb2 samples. When expressing recombinant gene *H5^Indo^*, activation of the UPR also correlated with the initiation of HA protein accumulation in the biomass between 1 and 3 DPI (Figure 1B). This strongly suggests that the cellular machinery involved in ER protein folding and maturation started to be mobilized as soon as the production of H5 began. Considering that this protein accumulates to levels that sustain the commercial production of influenza vaccine candidates (D’Aoust *et al.,* 2008; Landry *et al.,* 2010), and that the process results in somewhat mild stress symptoms on the plants (Figure 1A; Hamel *et al.,* 2023), early and transient activation of the UPR was apparently sufficient to cope with increased cellular needs associated to foreign HA protein secretion and to prevent deleterious effects that would have been caused by an unresolved ER stress. It is also important to keep in mind that despite reduced expression of UPR genes between 3 and 5 DPI in H5 samples, proteomics confirms that the level of several UPR proteins was still enhanced at 6 DPI (Table S1; Hamel *et al.,* 2023). Cellular benefits from a transient activation of the UPR thus appear to last for a longer timeframe, at least during expression of the foreign protein H5.

### Conservation of UPR signaling branches in N. benthamiana

For H5 and mAb2 samples, in which activation of the UPR was the strongest, one of the most striking observations is the fact that expression patterns for most UPR genes closely mirrored the expression patterns of *NbbZIP60s* (Figure 2C). Since unconventional splicing of *bZIP60* transcripts reflects IRE1 activity, this strongly suggests that this branch of the UPR is involved in foreign protein expression and that UPR genes examined are direct genetic targets of *NbbZIP60*. In plants, the UPR also comprises a second branch, which is initiated by BiP dissociation from ER membrane-tethered TFs bZIP17 and bZIP28 (Liu and Howell, 2010; Iwata and Koizumi, 2012; Duwi Fanata *et al.,* 2013). bZIPs can then cruise along the endomembrane system and end up in the Golgi apparatus, where clipping mediated by Golgi-associated proteases allows for their release from anchoring membrane. Free TFs then translocate to the nucleus, where they upregulate the expression of UPR genes in conjunction with bZIP60. Using protein sequences of Arabidopsis bZIP17 and bZIP28 as queries, blast of the *N. benthamiana* genome identified three genes encoding proteins that share about 50% identify with their respective homologs in Arabidopsis (Figure S5A). Although relatively low overall, sequence identity of the protein products from Niben101Scf32851g00038 (herein termed *NbbZIP17*), Niben101Scf03647g01004 (*NbbZIP28a*), and Niben101Scf00077g08013 (*NbbZIP28b*) is much higher within functional regions, including the bZIP domain, the TMD, and clipping sites targeted by Golgi-associated proteases (Figure S5B). Since the promoter region of every UPR gene examined here contains at least one *cis* regulatory element involved in UPR signaling (Figure S6), these TFs likely function in conjugation with NbbZIP60 to promote the expression of UPR genes during foreign protein expression. The development of an assay to monitor NbbZIP17 and NbbZIP28 clipping would however be required to formally confirm this hypothesis.

### Activation of the UPR and plant defense gene expression

Considering that expression of influenza HA induced a transient activation of the UPR, we also hypothesize that necrotic symptoms observed on leaves expressing this product were more likely caused by activation of plant immunity. At 6 DPI, H5 expression was shown to induce a unique molecular signature, including strong and specific upregulation of genes involved in oxylipin signaling and responses, as well as in the activation of oxidative stress responses (Hamel *et al.,* 2023). As many of these genes typically respond to wounding and herbivory, their upregulation was likely associated with budding of the VLPs, which highjack lipids from the PM of plant cells. At the molecular level, these multiple membrane budding events may be perceived as “micro-wounds” that would induce plant immunity. While this portrait was drawn using biomass harvested at a single time point after 6 days of expression, the time course approach employed here confirmed that upregulation of these defense genes comes after the peak of UPR activation at 3 DPI. Immunoblotting also indicated that defense gene upregulation correlated with a high increase in H5 protein accumulation (Figure 1B). Taken as a whole, these findings suggest that early activation of the UPR favors high production of the HA protein, in turn leading to the activation of plant immunity and eventually in the onset of leaf necrosis. This view is also supported by the fact that spraying of the leaves with an antioxidant solution of ascorbic acid helps to reduce HA-induced defenses and development of leaf necrotic symptoms, with no negative impact on foreign protein accumulation *in planta* (Hamel *et al.,* 2023).

In the case of mAb2 samples, in which the strongest necrotic symptoms were induced, expression from most defense genes examined was also induced compared to controls. When compared to H5 samples, induction of these defense genes was however of lower intensity and delayed by several days. As a result, late upregulation of defense genes was perhaps caused by the fact that at this stage of expression, plant cells were already dysfunctional and on the verge of collapsing. On the opposite, expression of UPR genes was induced early at 3 DPI, despite undetectable level of the mAb2 product until 7 DPI (Figure 1E). High UPR gene expression was then sustained up to end of leaf sampling at 9 DPI, suggesting that induced countermeasures never solved the issue associated to the expression of this antibody. In turn, this led to poor accumulation of the recombinant product and to strong plant cell death activation. While leaves expressing HA and mAb2 products both displayed necrotic symptoms, signaling pathways that induced this response were seemingly different, likely explaining why cell death lesions from the two conditions were macroscopically dissimilar (Figure 1A).

### Optimization of molecular farming approaches using components of the UPR

Achieving high recombinant protein yields *in planta* is one of the key aspects to ensure the success and sustainability of plant molecular farming approaches. While the expression of many foreign proteins is currently performed using wild-type plants, editing of the host plant genome can be envisioned as a way to optimize productivity or product quality. Alternatively, the co-expression of helper proteins can lead to an increase in foreign protein yields, as shown in *N. benthamiana* with the use of protease inhibitors that favor the accumulation of recombinant antibodies (Goulet *et al.,* 2012; Robert *et al.,* 2013; Jutras *et al.,* 2016; Grosse-Holz *et al.,* 2018). As for UPR proteins, heterologous expression of a human CRT was shown to improve production of viral glycoproteins that initially showed poor accumulation levels *in planta* (Margolin *et al.,* 2020). The results presented here highlight a series of *N. benthamiana* genes that could be used as markers of UPR activation during a stress response, or expression of foreign proteins *in planta*. For complex or even refractory products, the UPR genes identified here may also constitute an interesting list of potential endogenous protein helpers to be co-expressed as a way to enhance ERQC functions of plant cell biofactories, or to prevent undesirable effects provoked by an unresolved ER stress during foreign protein expression.

## Materials and methods

### Seed germination and plant growth

Seeds of *N. benthamiana* were spread on pre-wetted peat mix plugs (Ellepot) and placed in a germination chamber for 2 days, where conditions were as follows: 28°C/28°C day/night temperature, 16 h photoperiod, 90% relative humidity, and light intensity of 7 µmol m^-2^ s^-1^. Germinated plantlets were next transferred in a growth chamber for 15 days, where conditions were as follows: mean temperature of 28°C over 24 h, 16 h photoperiod, mean relative humidity of 66% over 24 h, 800 ppm carbon dioxide (CO_2_) injected only during the photo-phase, and light intensity of 150 µmol m^-2^ s^-1^. During this time, watering and fertilization were provided as needed. After 2 weeks, peat mix plugs were transferred to 4 inches pots containing pre-wetted peat-based soil mix (Agro-Mix). Freshly transferred plantlets were then moved to a greenhouse, where conditions were as follows: mean temperature of 25°C over 24 h, 16 h photoperiod, mean relative humidity of 66% over 24 h, 800 to 1,000 ppm CO_2_ injected only during the photo-phase, and light intensity according to natural conditions, but supplemented with artificial high pressure sodium lights at 160 µmol m^-2^ s^-1^. In the greenhouse, watering and fertilization were provided as needed. Growth was allowed to proceed for an average of 20 additional days, until the plants were ready for agroinfiltration.

### Binary vector constructs

For VLP expression, sequences from the mature HA protein of pandemic influenza virus strain H5 Indonesia (H5/A/Indonesia/05/2005; H5^Indo^) were fused to the signal peptide of a *Medicago sativa* (alfalfa) PDI using PCR-based methods. Once assembled, the chimeric *H5^Indo^* gene was reamplified by PCR and then introduced in the T-DNA region of a pCAMBIA binary vector previously linearized with restriction enzymes *SacII* and *StuI* using the In-Fusion cloning system (Clontech). The same methods and PDI signal peptide were used to clone antibody chain genes, which allowed expression of recombinant antibodies mAb1 and mAb2. Expression of recombinant genes *H5^Indo^*, *mAb LC* and *mAb HC* was driven by a 2X35S promoter from the cauliflower mosaic virus (CaMV). Expression cassettes also comprised 5’- and 3’-untranslated regions (UTRs) from the cowpea mosaic virus (CPMV), and the *Agrobacterium nopaline synthase* (*NOS*) gene terminator. To prevent silencing induced by recombinant gene expression *in planta*, the T-DNA region of binary vectors also included the suppressor of RNA silencing gene *P19*, under the control of a plastocyanin promoter and terminator. For P19 samples, a binary vector allowing expression of P19 only was employed.

### Agrobacterium cultures and plant infiltration

Binary vectors were transformed by heat shock in *Agrobacterium* strain AGL1. Transformed bacteria were plated on Luria-Bertani (LB) medium, with appropriate antibiotics selection (kanamycin 50 µg/ml). Colonies were allowed to develop at 28°C for 2 days. Using isolated colonies, frozen glycerol stocks were prepared and placed at −80°C for long term storage. When ready, frozen bacterial stocks were thawed at room temperature before transfer in pre-culture shake flasks containing LB medium with antibiotics selection (kanamycin 50 µg/ml). Bacterial pre-cultures were grown for 18 h at 28°C with shaking at 200 rpm. While keeping kanamycin selection, pre-cultures were transferred to larger shake flasks and bacteria were allowed to develop for an extra 18 h at 28°C with shaking at 200 rpm. Using a spectrophotometer (Implen), bacterial inoculums were prepared by diluting appropriate volumes of the bacterial cultures in resuspension buffer (10 mM MgCl_2_, 5 mM MES, pH 5.6). A final OD_600_ of 0.6 was employed for all experiments and vacuum infiltration was performed by submerging whole plant shoots in the appropriate bacterial suspension.

### Transient protein expression and biomass harvesting

Recombinant protein accumulation was allowed to proceed for several days, as indicated. For all experiments, expression took place in condition-controlled plant growth chambers, where settings were as follows: 20°C/20°C day/night temperature, 16 h photoperiod, 80% relative humidity, and light intensity of 150 µmol m^-2^ s^-1^. Watering was performed every other day, with no fertilizer supplied during the expression phase. For biomass harvesting, leaves of similar developmental stage were selected using the leaf plastochron index (Meicenheimer, 2014). The fourth and fifth fully expanded leaves starting from the top of each plant were harvested without petiole. Freshly cut leaves were placed in pre-frozen 50 mL Falcon tubes, before flash freezing in liquid nitrogen. Frozen biomass was stored at −80°C until ready for analysis. Using pre-chilled mortars and pestles, foliar tissue was ground and homogenized into powder using liquid nitrogen. Each sample was made from four leaves collected on two randomly selected plants. The average results presented were obtained from at least three biological replicates.

### Protein extraction and quantification

For protein extraction, 1 g of frozen biomass powder was taken out of the −80°C freezer and placed on ice. A 2 mL volume of extraction buffer (50 mM Tris, 500 mM NaCl, pH 8.0) was added, followed by 20 µL of 100 mM phenylmethanesulfonyl fluoride (PMSF) and 2 µL of 0.4 g/mL metabisulfite. Quickly after addition of all solutions, samples were crushed for 45 sec using a Polytron homogenizer (ULTRA-TURRAX^®^ T25 basic) at maximum speed. One mL of each sample was transferred to a prechilled eppendorf tube and centrifuged at 10 000 x g for 10 min at 4°C. Supernatants were carefully recovered, transferred to new eppendorf tubes, and kept on ice until determination of protein concentration. To quantify protein content from crude extracts, the Bradford method was employed, with bovine serum albumin as a protein standard.

### Western blotting, HMG assays, and gel densitometry

For western blotting, total protein extracts were diluted in extraction buffer and mixed with 5X Laemmli sample loading buffer to reach a final concentration of 0,5 µg/µL. Protein samples were denatured at 95°C for 5 min, followed by a quick spin using a microcentrifuge. 20 µL of each denatured protein extract (10 µg) was then loaded on Criterion^TM^ XT Precast polyacrylamide gels 4-12% Bis-Tris and separated at 110 volts for 105 min. Using transfer buffer (25 mM Tris, 192 mM Glycine, 10% methanol), proteins were next electrotransferred onto a polyvinylidene difluoride (PVDF) membrane at 100 volts. After 30 min, membranes were placed in blocking solution: 1X Tris-Buffered Saline with Tween-20 (TBS-T; 50 mM Tris, pH 7.5, 150 mM NaCl, 0,1% (v/v) Tween-20), with 5% nonfat dried milk. Membranes were blocked overnight at 4°C with gentle shaking. The next morning, blocking solution was removed and primary antibodies were incubated at room temperature for 60 min with gentle shaking in 1X TBS-T, 2% nonfat dried milk solution. After 4 washes in 1X TBS-T, secondary antibodies were added and incubated at room temperature for 60 min with gentle shaking in 1X TBS-T, 2% nonfat dried milk solution. After 4 extra washes in 1X TBS-T, Luminata^TM^ Western HRP Chemiluminescence Substrate (Thermo Fisher Scientific) was added to the membranes and protein complexes were visualized under the chemiluminescence mode of an Imager 600 apparatus (Amersham). Antibody dilutions were as follows: anti-HA A/Indonesia/05/2005 (H5N1; CBER): 1/5,000 (primary antibody). Rabbit anti-sheep (JIR): 1/10,000 (secondary antibody).

For HMG assays, turkey red blood cells were diluted to a concentration of 0,25% (v/v) in phosphate-buffered saline solution (PBS; 0.1 M PO_4_, 0.15 M NaCl, pH 7.2). While keeping red blood cells on ice, protein samples were diluted in extraction buffer using 1/384 and 1/576 ratios. For each dilution, 200 µL of total protein extract was transferred to the first row of a 96-well plate. Eight serial dilutions were next performed using 100 µL of protein extract mixed to 100 µL of PBS buffer previously poured in each plate well. Following serial dilutions, 100 µL of the red blood cell solution was added to protein extracts. After thorough mixing, samples were incubated overnight at room temperature. HA activity was scored visually on the next day.

For gel densitometry, total protein extracts were diluted in extraction buffer and mixed with 5X Laemmli sample loading buffer. The protein samples were denatured at 95°C for 5 min, followed by a quick spin using a microcentrifuge. For each sample, 2 and 4 µg of denatured protein extracts were loaded on Criterion^TM^ XT Precast polyacrylamide gels 4-12% Bis-Tris. To quantify fully assembled antibodies, a standard curve consisting of 1.5 µg to 0.125 µg of purified IgG1 (Sigma) was added on each gel. Protein separation was done at 110 volts for 105 min and staining of the gels was done with Coomassie blue.

### RNA extractions and RNA quantification

Using the RNeasy commercial kit (Qiagen), 100 mg of frozen biomass powder was used for RNA extractions. Residual DNA was removed using the RNase-free DNase Set (Qiagen). Concentration of RNA extracts was determined using a spectrophotometer (Implen) and integrity evaluated using a 2100 BioAnalyzer (Agilent). For long term storage, RNA extracts were stabilized by adding the RNAseOUT recombinant ribonuclease inhibitor (Thermo Fisher Scientific), before freezing at −80°C until further analysis.

### RTqPCR analyses

For each sample, 1 µg of RNA was reverse transcribed into cDNA using the QuantiTect Reverse Transcription Kit (Qiagen). Transcript quantification was performed in 96-well plates, using the ABI PRISM 7500 Fast real-time PCR system and custom data analysis software (Thermo Fisher Scientific). Each reaction contained the equivalent of 5 ng cDNA as a template, 0.5 µM of forward and reverse primers, and 1X QuantiTect SYBR Green Master Mix (Qiagen) for a total reaction volume of 10 µL. RTqPCR runs were done under the SYBR Green amplification mode and cycling conditions were as follows: 15 min incubation at 95°C, followed by 40 amplification cycles at 95°C for 5 sec, 60°C for 30 sec, and 65°C for 90 sec. Reactions in the absence of cDNA template were conducted as negative controls and melting curve analyses were performed to confirm the lack of primer dimer formation and amplification specificity. Resulting fluorescence and cycle threshold (Ct) values were next exported to the Microsoft Excel software. To correct for biological variability and technical variations during RNA extraction, quantification, or reverse transcription, expression from six housekeeping genes (*NbACT1*, *NbVATP*, *NbSAND*, *NbUBQ1*, *NbEF1-*α, and *NbGAPDH1*) was used to normalize expression data (Vandesompele *et al.,* 2002; Hamel *et al.,* 2023). Normalized numbers of molecules per ng of RNA were deduced using the 2^-ΔΔCt^ method (Livak and Schmittgen, 2001; Bustin *et al.,* 2009) and standard curves derived from known quantities of phage lambda DNA. Standard deviation related to the within-treatment biological variation was calculated in accordance with the error propagation rules. Sequences of all primers used in this study are available in Table S3.

### Phylogenetic analyses

For each protein family to be analyzed, the genome of *N. benthamiana* (https://solgenomics.net/) was searched using full-length amino acid sequences of Arabidopsis homologs as queries. Full-length protein sequences retrieved from blast analyses were next aligned with ClustalW. Alignment parameters were as follows: for pairwise alignments, 10.0 for gap opening and 0.1 for gap extension. Parameters for multiple alignments were 10.0 for gap opening and 0,20 for gap extension. The resulting alignments were submitted to the MEGA5 software and neighbor-joining trees derived from 5,000 replicates were generated. Bootstrap values are indicated on the node of each branch. More details on phylogenetic analyses are provided in the legend of relevant figures.

### Statistical analyses

Statistical analyses of RTqPCR data were performed on Graph Pad Prism 9.5.0, using two-way ANOVA with time points as row factors and conditions as column factors. Conditions were then analysed by a post-hoc Tukey’s multiple comparison test with an alpha threshold of 0.05. For clarity and to limit the number of statistical groups generated, control conditions (NI and Mock) were omitted from statistical analyses, which were thus only performed on agroinfiltrated conditions (P19, H5, mAb1, and mAb2). Groups are labeled with a compact letter display. Groups that do not share the same letter(s) are statistically different. Complete results from statistical analyses can be found in Table S4.

### Search for cis regulatory elements

To locate *cis* regulatory elements within the promoter of UPR genes, the putative promoter regions of each gene tested were retrieved from the genome sequences of *N. benthamiana* (https://solgenomics.net/). For the analysis, a 1,000 bp region located upstream of the annotated start codon was employed. Motif search was performed in both sequence orientations using the Find Individual Motif Occurrences (FIMO) software (Grant *et al.,* 2011). As they were previously involved in the UPR (Iwata and Koizumi, 2005; Iwata *et al.,* 2008; Liu and Howell, 2016), *cis* regulatory elements that were examined were as follows: PUPRE (ATTGGTCCACGTCATC), ERSE (CCAATN_10_CACG), ERSE2 (ATTGGN_2_CACG), UPRE2 (GATGACGCGTAC), UPRE3 (TCATCG), and XBP-BS (GATGACGTGK).

## Acknowledgments and funding

This work would not have been possible without the support of the Biomass Production and Research & Innovation teams of Medicago. Their contribution includes, but is not limited to, cloning of genetic constructs, inoculum preparation, growth of the plants and agroinfiltration. Authors also wish to acknowledge Gervais Pelletier and Dr. Armand Séguin from Natural Resources Canada for technical support and advice on RTqPCR. For support during the proteomics study, authors are also grateful to Antoine Leuthreau, graduate student at the Laval University Plant Science Department. Funding for this work was provided by Medicago Inc. This work was also supported by a Collaborative Research and Development grant from the Natural Sciences and Engineering Research Council of Canada to Dominique Michaud (CRDPJ 495852).

## Author contributions

L.P.H., D.M., and M.A.D. designed the research, supervised the project, and analyzed the data. L.P.H. and M.A.C. assembled the figures and drafted the manuscript. P.O.L. managed the production of genetic constructs used to express influenza HA and antibodies. M.C.G. contributed to the proteomics study. L.P.H., F.P.G., and M.A.C. managed data and performed searches within RNAseq and proteomics data. R.T., F.P.G., M.E.P., and M.A.C. performed RTqPCR, assays for recombinant protein quantification and western blots. F.P.G. and M.A.C. performed the statistical analyses. M.A.C. performed searches for *cis* regulatory elements. All authors read, helped to edit, and approved the final version of the manuscript.

## Data availability statement

All data discussed in this study can be found in the manuscript and in the Supplementary Materials.

## Supporting information

Additional supporting information may be found online in the Supporting Information section at the end of the article.

**Table S1.** UPR proteins upregulated by P19 expression or co-expression of P19 and influenza HA.

**Table S2.** Name and identification number of UPR genes from *A. thaliana* and *N. benthamiana*.

**Table S3.** RTqPCR primers used in this study.

**Table S4.** Statistical analyses of the RTqPCR results.

## Figure legends

**Figure S1.**
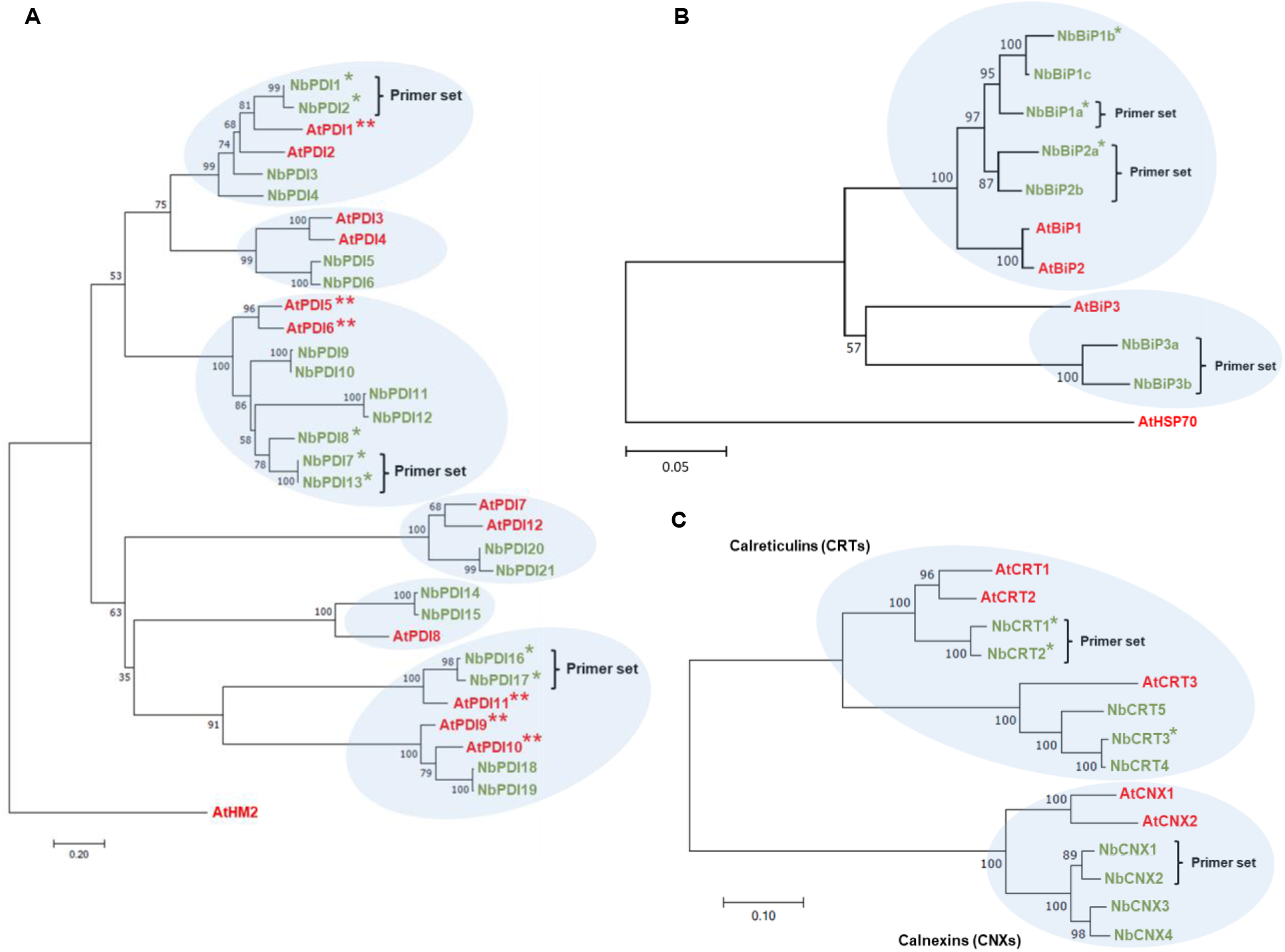
Phylogeny of *N. benthamiana* PDIs and ER-resident chaperones. (A) The genome of *N. benthamiana* was searched using full-length amino acid sequences of AtPDI1 (At3g54960) as a query. Full-length protein sequences that were retrieved were then aligned with ClustalW, using thioredoxin AtHM2 (At4g03520) as an outgroup. The resulting alignments were submitted to the MEGA5 software, and a neighbor-joining tree derived from 5,000 replicates was generated. Bootstrap values are indicated on the node of each branch. (B) For the phylogeny of BiPs, the same approach as above was employed, except that AtBiP1 (At5g28540) was used as a query to blast the genome and AtHSP70 (At3g12580) served as an outgroup. (C) For the phylogeny of CRT and CNX lectins, the same approach as above was employed, except that AtCRT1 (At1g56340) and AtCNX1 (At5g61790) were used as queries to blast the genome. The CNX cluster served as an outgroup for CRTs, and vice versa. *A. thaliana* (At) proteins are shown in red, while *N. benthamiana* (Nb) proteins are shown in green. Brackets highlight genes targeted by RTqPCR analyses. Single asterisks (*) highlight proteins identified by proteomics (Table S1; Hamel *et al.,* 2023). Double asterisks (**) denote AtPDIs involved in the UPR (Lu and Christopher, 2008). Gene model number corresponding to Arabidopsis and *N. benthamiana* proteins can be found in Table S2.

**Figure S2.**
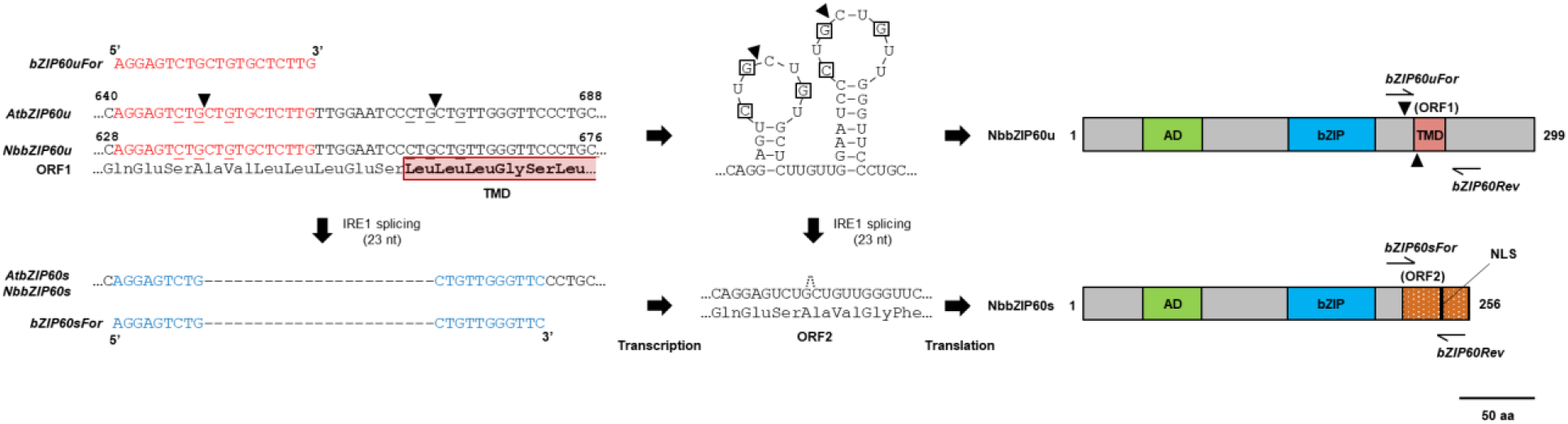
Primer design to assess unconventional splicing of *NbbZIP60*. Left section: nucleotide (nt) sequence alignment confirming conservation of unconventional splicing regions within *bZIP60* transcripts from *A. thaliana* (*AtbZIP60u*) and *N. benthamiana* (*NbbZIP60u*). Three-letter amino acid (aa) code of the corresponding open reading frame (ORF1) is shown and start of the transmembrane domain (TMD) is boxed in red. Conserved nt from the repeated CXGXXG motifs are underlined. As part of the stem-loop structures recognized by IRE1, these nt are essential for mRNA splicing. Black triangles indicate predicted mRNA splicing sites. After removal of a 23 nt intron, sequence of spliced *bZIP60* (*bZIP60s*) is shown. Sequences from forward (For) primers used to discriminate spliced and unspliced versions of the transcripts are shown in blue (*bZIP60sFor*) and red (*bZIP60uFor*), respectively. Middle section: predicted stem-loops from the *bZIP60u* mRNA. Conserved nt from the repeated CXGXXG motifs are boxed and black triangles highlight predicted mRNA splicing sites. After removal of the 23 nt intron by IRE1, nt and deduced aa sequences from the new open reading frame that is created (ORF2) are shown. Right section: at-scale topology from the predicted NbbZIP60u and NbbZIP60s proteins. The transcriptional activation domain (AD) is shown in green. The basic leucine zipper (bZIP) domain, which is required for protein dimerization and DNA-binding, is shown in blue. The TMD of NbbZIP60u is shown in red and the alternative C-terminus region from NbbZIP60s in textured orange. The TMD is no longer translated, but this region now includes a putative nuclear localization signal (NLS) depicted in black. Based on corresponding transcripts sequences, relative position of the For and reverse (Rev) primers used for RTqPCR are shown. The Rev primer was the same for both transcripts (*bZIP60Rev*). Adapted from Nagashima *et al.,* 2011.

**Figure S3.**
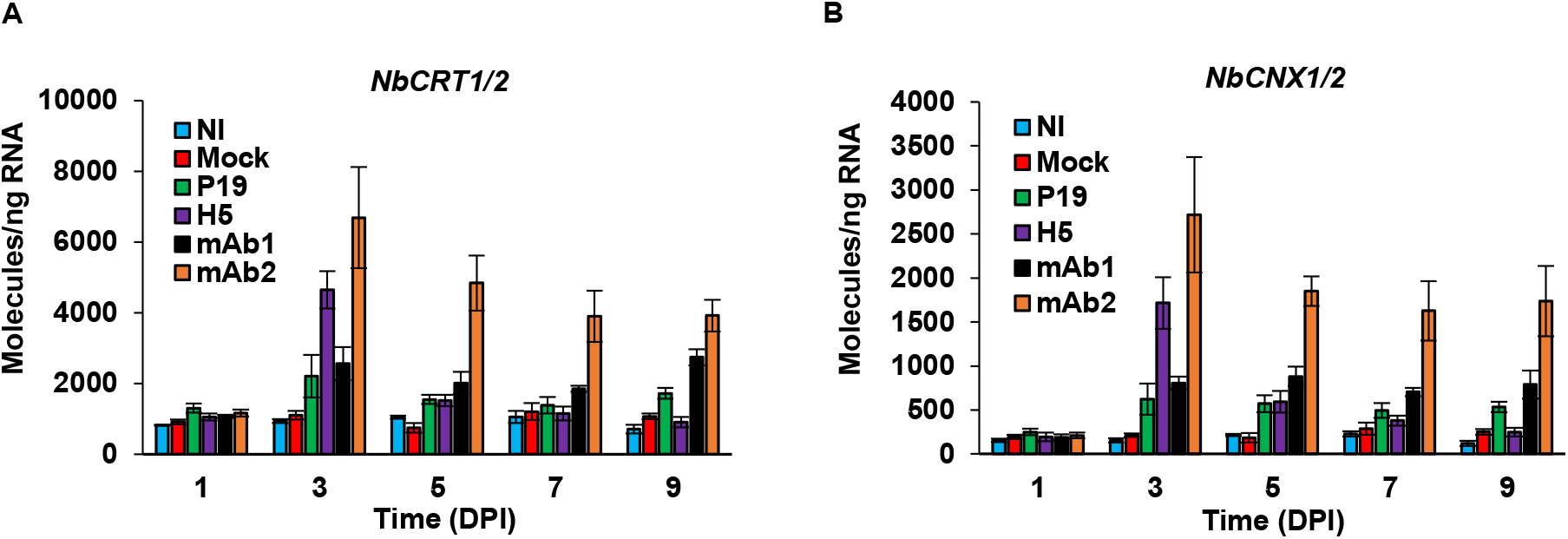
Expression of ERQC genes: *CRTs* and *CNXs*. Expression of genes encoding for ER resident chaperones of the CRT (A) and CNX (B) families, as measured by RTqPCR. For each time point in days post-infiltration (DPI), results are expressed in numbers of molecules per ng of RNA. Condition names are as follows: NI, non-infiltrated leaves; Mock, leaves infiltrated with buffer only; P19, leaves infiltrated with AGL1 and expressing P19 only; H5, leaves infiltrated with AGL1 and co-expressing P19 and H5^Indo^; mAb1 (and mAb2), leaves infiltrated with AGL1 and co-expressing P19, monoclonal antibody mAb1 (or mAb2) light (LC) and heavy (HC) chains.

**Figure S4.**
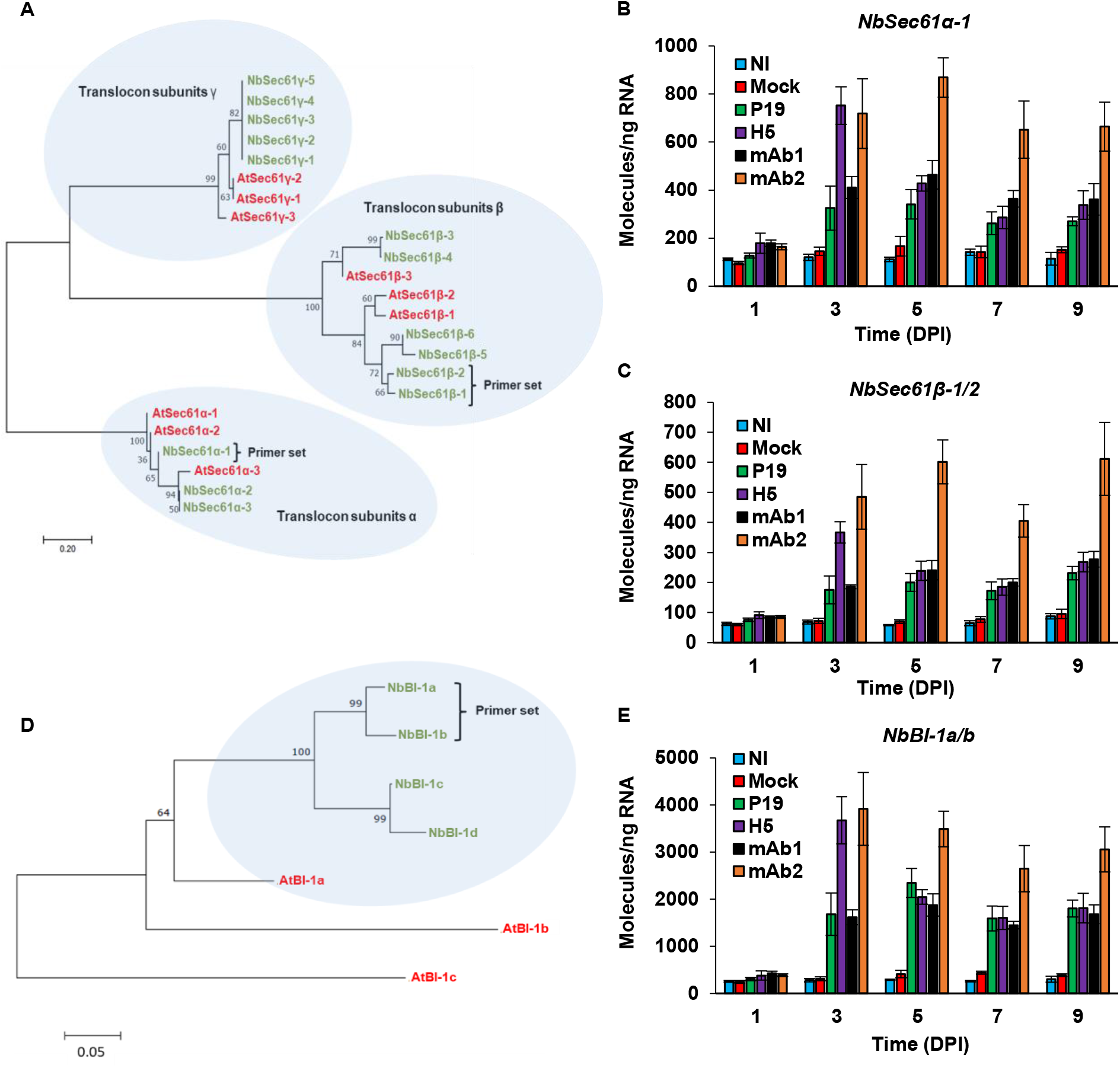
Phylogeny of other UPR proteins and expression of some of their corresponding genes. (A) For Sec61 proteins, the genome of *N. benthamiana* was searched using full-length amino acid sequences of AtSec61α-1 (At1g29310), AtSec61β-1 (At2g45070), and AtSec61γ-1 (Atg424920) as queries. Retrieved full-length protein sequences were then aligned with ClustalW. The distinct Sec61 protein clusters served as outgroups for each other. The resulting alignments were submitted to the MEGA5 software, and a neighbor-joining tree derived from 5,000 replicates was generated. Bootstrap values are indicated on the node of each branch. *A. thaliana* (At) proteins are shown in red, while *N. benthamiana* (Nb) proteins are shown in green. Brackets highlight genes targeted by RTqPCR analyses. Gene model number corresponding to Arabidopsis and *N. benthamiana* proteins can be found in Table S2. Expression of genes encoding for Sec61 proteins NbSec61α-1 (B) as well as NbSec61β-1 and NbSec61β-2 (C), as measured by RTqPCR. For each time point in days post-infiltration (DPI), results are expressed in numbers of molecules per ng of RNA. (D) For the phylogeny of BI-1 proteins, the same approach as above was employed except that AtBI-1a (AT5G47120) was used as a query to blast the genome. Distant homolog AtBI-1c (At5g47130) served as an outgroup. Expression of genes encoding for BI-1 proteins NbBI-1a and NbBI-1b (E), as measured by RTqPCR. Condition names are as follows: NI, non-infiltrated leaves; Mock, leaves infiltrated with buffer only; P19, leaves infiltrated with AGL1 and expressing P19 only; H5, leaves infiltrated with AGL1 and co-expressing P19 and H5^Indo^; mAb1 (and mAb2), leaves infiltrated with AGL1 and co-expressing P19, monoclonal antibody mAb1 (or mAb2) light (LC) and heavy (HC) chains.

**Figure S5.**
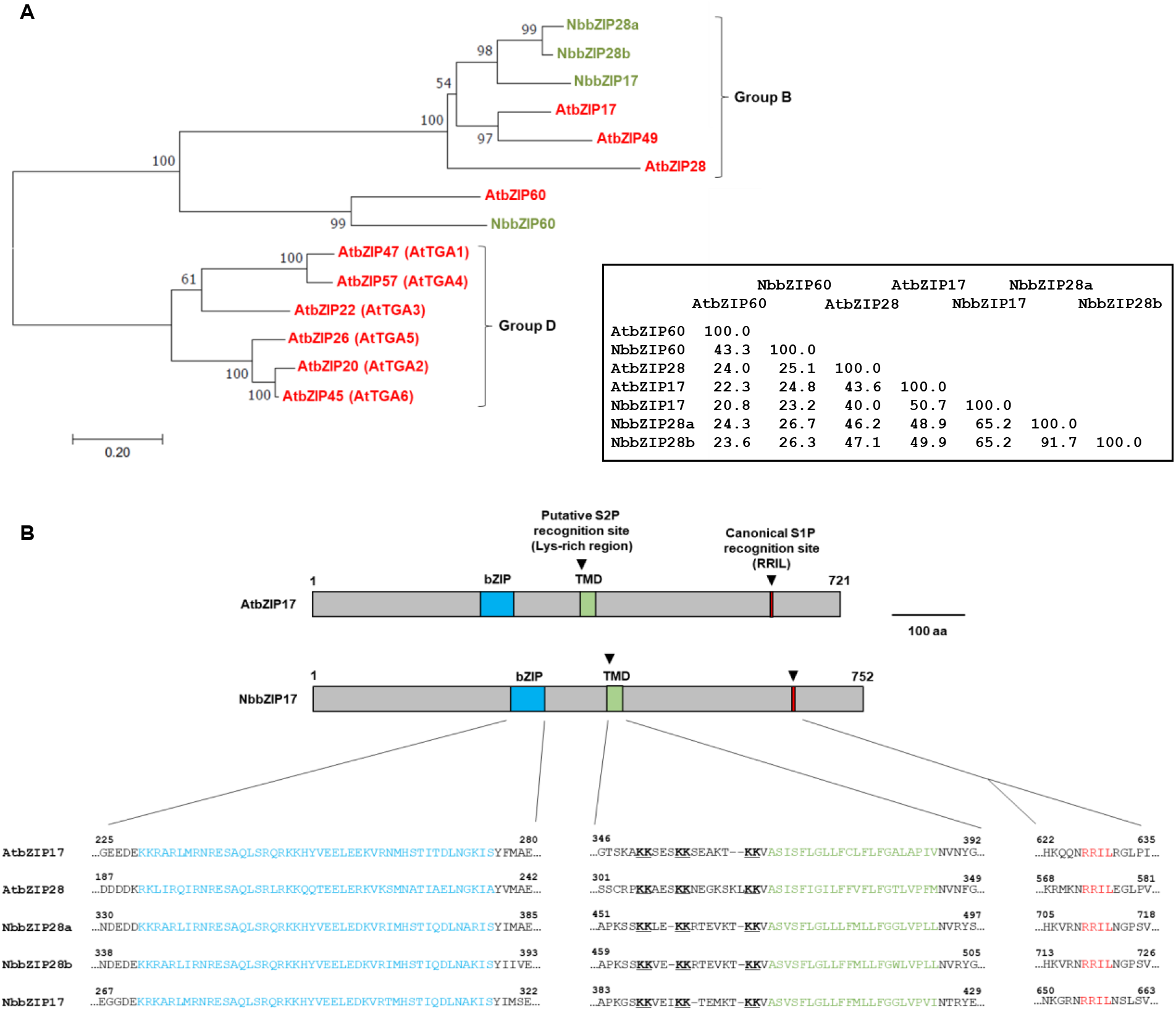
Phylogeny of UPR activating bZIPs and conservation of bZIP17 and bZIP28 in *N. benthamiana*. (A) The genome of *N. benthamiana* was searched using full-length amino acid sequences of AtbZIP17 (At2g40950), AtbZIP28 (At3g10800), and AtbZIP60 (At1g42990) as queries. Full-length protein sequences that were retrieved were then aligned with ClustalW, using a set of defense-related bZIPs from Arabidopsis as an outgroup (Group D bZIPs; Stotz *et al.,* 2013). Resulting alignments were submitted to the MEGA5 software and a neighbor-joining tree derived from 5,000 replicates was generated. Bootstrap values are indicated on the node of each branch. *A. thaliana* (At) proteins are shown in red, while *N. benthamiana* (Nb) proteins are shown in green. Brackets highlight bZIP groups, as defined previously in Arabidopsis (Jakoby *et al.,* 2002). In this classification, bZIP60s remain unclassified. For UPR activating bZIPs, a percent identity matrix is shown on the right. Gene model number corresponding to bZIP17, bZIP28, and bZIP60 homologs from Arabidopsis and *N. benthamiana* can be found in Table S2. (B) At-scale topology of proteins AtbZIP17 and NbbZIP17. The basic leucine zipper (bZIP) domain, which is required for protein dimerization and DNA-binding, is shown in blue. The transmembrane domain (TMD) is shown in green. The canonical recognition site of Golgi-associated protease S1P (RXXL motif) is shown in red. Clipping site of S1P and predicted clipping site of the Golgi-associated protease S2P are depicted by black triangles. Conservation of functional regions within bZIP17 and bZIP28 homologs from Arabidopsis and *N. benthamiana* is depicted by a protein sequence alignment, with amino acids displayed using their one-letter code. S2P-mediated clipping is thought to occur within a lysine (Lys)-rich region located before the TMD and conserved Lys residues are bolded and underlined.

**Figure S6.**
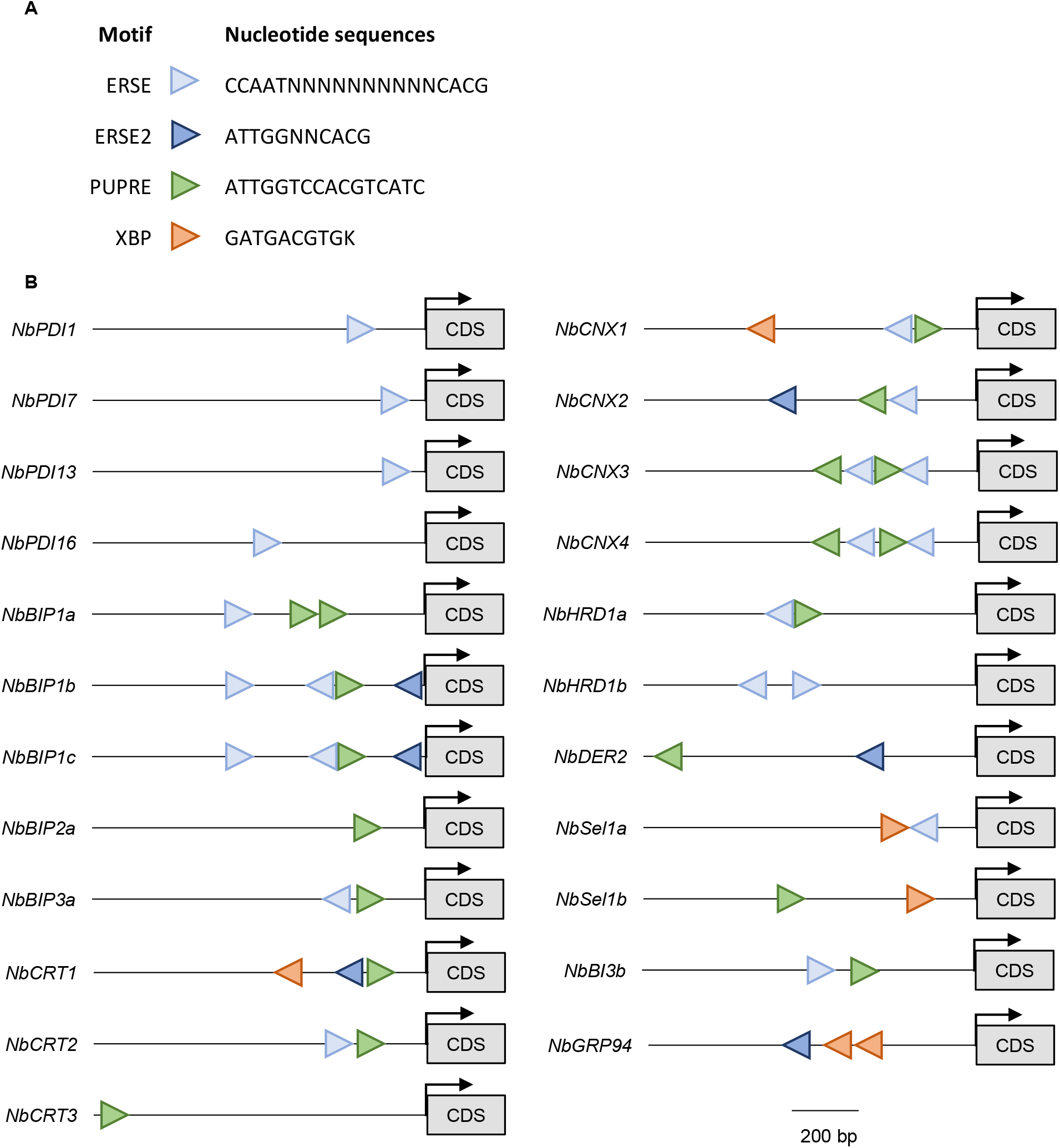
*Cis* regulatory elements in the promoter of UPR genes. (A) Consensus nucleotide sequences of *cis* regulatory elements that were examined. These motifs have previously been linked to UPR activation in plants (Iwata and Koizumi, 2005; Iwata *et al.,* 2008; Liu and Howell, 2016). Each motif is depicted by a color-coded triangle. "N" letters represent any nucleotide, while the "K" letter represents a guanine (G) or a thymine (T). (B) Using genome sequences from *N. benthamiana*, putative promoter regions from selected UPR genes were retrieved. For each gene, a 1000 base pairs (bp) region located upstream of the annotated start codon was employed and motif scanning was performed in both sequence orientations using the FIMO software (Grant *et al.,* 2011). Approximate location and orientation of each motif are indicated by color-coded triangles. Predicted start codon of each coding sequence (CDS) is highlighted by a black arrow.

